# Macrophage-targeted lipid nanoparticle delivery of microRNA-146a to mitigate hemorrhagic shock-induced acute respiratory distress syndrome

**DOI:** 10.1101/2023.02.17.529007

**Authors:** Qinqin Fei, Emily M. Shalosky, Ryelie Barnes, Vasudha C. Shukla, Megan N. Ballinger, Laszlo Farkas, Robert J. Lee, Samir N. Ghadiali, Joshua A. Englert

**Author notes:** **Corresponding Authors** Joshua A. Englert - Division of Pharmaceutics and Pharmacology, College of Pharmacy, The Ohio State University, 500 West 12th Avenue, Columbus, OH 43210, USA. Division of Pulmonary, Critical Care, and Sleep Medicine, Department of Internal Medicine, The Ohio State University Wexner Medical Center, 473 West 12^th^ Avenue, Columbus OH 43210, USA. The Davis Heart and Lung Research Institute, The Ohio State University Wexner Medical Center, 473 West 12th Avenue, Columbus, OH 43210;, Samir N. Ghadiali - Department of Biomedical Engineering, The Ohio State University, 140 West 19th Avenue, Columbus, OH 43210, USA. The Davis Heart and Lung Research Institute, The Ohio State University Wexner Medical Center, 473 West 12th Avenue, Columbus, OH 43210.

## Abstract

The pro-inflammatory response of alveolar macrophages to injurious physical forces during mechanical ventilation is regulated by the anti-inflammatory microRNA, miR-146a. Increasing miR-146a expression to supraphysiologic levels using untargeted lipid nanoparticles reduces ventilator-induced lung injury, but requires a high initial dose of miR-146a making it less clinically applicable. In this study, we developed mannosylated lipid nanoparticles that can effectively mitigate lung injury at the initiation of mechanical ventilation with lower doses of miR-146a. We used a physiologically relevant humanized *in vitro* co-culture system to evaluate the cell-specific targeting efficiency of the mannosylated lipid nanoparticle. We discovered that mannosylated lipid nanoparticles preferentially deliver miR-146a to alveolar macrophages and reduce force-induced inflammation *in vitro*. Our *in vivo* study using a clinically relevant mouse model of hemorrhagic shock-induced acute respiratory distress syndrome demonstrated that delivery of a low dose miR-146a (0.1 nmol) using mannosylated lipid nanoparticles dramatically increases miR-146a in mouse alveolar macrophages and decreases lung inflammation. These data suggest that mannosylated lipid nanoparticles may have therapeutic potential to mitigate lung injury during mechanical ventilation.

## Introduction

The acute respiratory distress syndrome (ARDS) is a life-threatening condition that occurs in patients with trauma, sepsis, pneumonia, as well as coronavirus disease 2019 (COVID-19)^1,2^. Patients with ARDS require life-support therapy with mechanical ventilation (MV)^3–5^; however, MV generates physical forces that damage the alveolar-capillary barrier, trigger the release of proinflammatory mediators, and exacerbate lung injury in a process known as ventilator-induced lung injury (VILI)^3–5^. A recent study in our laboratory demonstrated that alveolar macrophages (AMs), the resident immune cells of the lung, respond to injurious forces during MV by upregulating an anti-inflammatory microRNA, miR-146a^6^; however, this endogenous response is insufficient to mitigate VILI^6^. We demonstrated that overexpression of miR-146a *in vivo* using untargeted lipid nanoparticles (LNPs) mitigates cytokine release and barrier disruption following injurious ventilation^6^. In our prior study, two doses of miR-146a delivered by untargeted LNPs were required to have a therapeutic effect^6^. We hypothesized that targeted delivery of miR-146a to the primary immune cells of the lung (i.e. alveolar macrophages) would be more effective than global overexpression. Therefore, we sought to develop targeted LNPs that preferrentially deliver anti-inflammatory microRNAs to AMs.

Different types of nanoparticles can be used to target specific cell types for drug delivery. One method to achieve this is by surface modification of nanoparticles with targeting ligands^7^. Previous studies have used ligand-anchored nanoparticle delivery systems to deliver drugs to AMs using ligands that are highly expressed on the surface of alveolar macrophages^8,9^. The mannose receptor (CD206) is a transmembrane glycoprotein expressed on the cell surface of tissue macrophages including those in the lung^10, 11^. It plays a key role in the recognition, internalization, and clearance of pathogens^10, 12^. Mannosylated nanoparticles are taken up through mannose receptor-mediated endocytosis which consequently enhances drug delivery to AMs^13–16^. The mannose receptor does not have an intracellular signaling motif on its cytoplasmic tail, which limits the potential for inflammation in response to ligand engagement^17^, thus minimizing immunogenicity of mannose-conjugated nanoparticles.

In this study, we tested the hypothesis that mannosylated LNPs enhance the delivery of miR-146a to AMs and more potently mitigate lung injury during MV compared to untargeted LNPs. We demonstrate that the mannosylated LNPs formulated in this study efficiently and preferentially deliver miR-146a to AMs. This enhanced delivery mitigated lung injury during mechanical ventilation and allowed for the use of lower doses of miR-146a and thus increases the translational potential of this technology.

## Results

### Fabrication and characterization of mannosylated lipid nanoparticles

We fabricated three formulations of mannosylated lipid nanoparticles with varying amounts of 1,2-dipalmitoyl-sn-glycero-3-phospho((ethyl-1’,2’,3’-triazole)triethyleneglycolmannose) (PA-PEG3-M) lipid (2%, 4.85%, 9.3%). The MLNPs were prepared using the ethanol injection and sonication method (**Figure *1*a**). miR-MLNPs had a spherical morphology observed under Cryo-TEM (**Figure *1*b**). The particle size and zeta potential of different MLNP formulations are shown in **Table 1**. The average diameter of the empty 9.3-MLNP as determined by dynamic light scattering was 189.9 ± 13.5 nm (mean ± SEM) with a polydispersity index (PDI) of 0.140 ± 0.020 (mean ± SEM) (**Figure *1*c** and **Table 1**). The miR-146a loaded 9.3-MLNP had a similar average diameter (218.7 ± 13.2 nm) and PDI (0.181 ± 0.033 nm) (**Figure *1*c** and **Table 1**). The average particle size of all MLNP formulations was less than 220 nm (**Table 1**) which is considered a suitable size for pulmonary gene delivery to the distal lung^18–20^. The size distribution of all the formulations was narrow as reflected by the low PDI (< 0.3) (**Table 1**). The zeta potential of all three formulations of empty MLNPs in distilled water (pH 6.5) was negative because of the negatively charged lipids. The zeta potential of the lipid nanoparticles shifted from negative to positive after loading of positively charged PEI/miR polyplex (**Table 1**). The stability of the MLNPs over time was evaluated by analyzing the particle size and PDI at days 0, 3, 7, and 14 after fabrication. The particle size of both empty and miR-146a loaded MLNPs did not change after 14 days of storage at 4°C, and PDI remained low (<0.3) (**Figure 1d**). Before testing the ability of MLNPs to deliver microRNAs to macrophages, we investigated whether the THP-1 human monocytic cell line and our human primary AMs express the mannose receptor in culture. Using immunofluorescence staining (**Figure S1a**) and western blotting (**Figure S1b-c**) it was found that primary AMs from multiple donors expressed the mannose receptor at high levels (**Figure S1b-c**). In contrast, THP-1 cells had low levels of CD-206 expression. Therefore, only primary AMs were used for subsequent studies.

**Figure 1.**
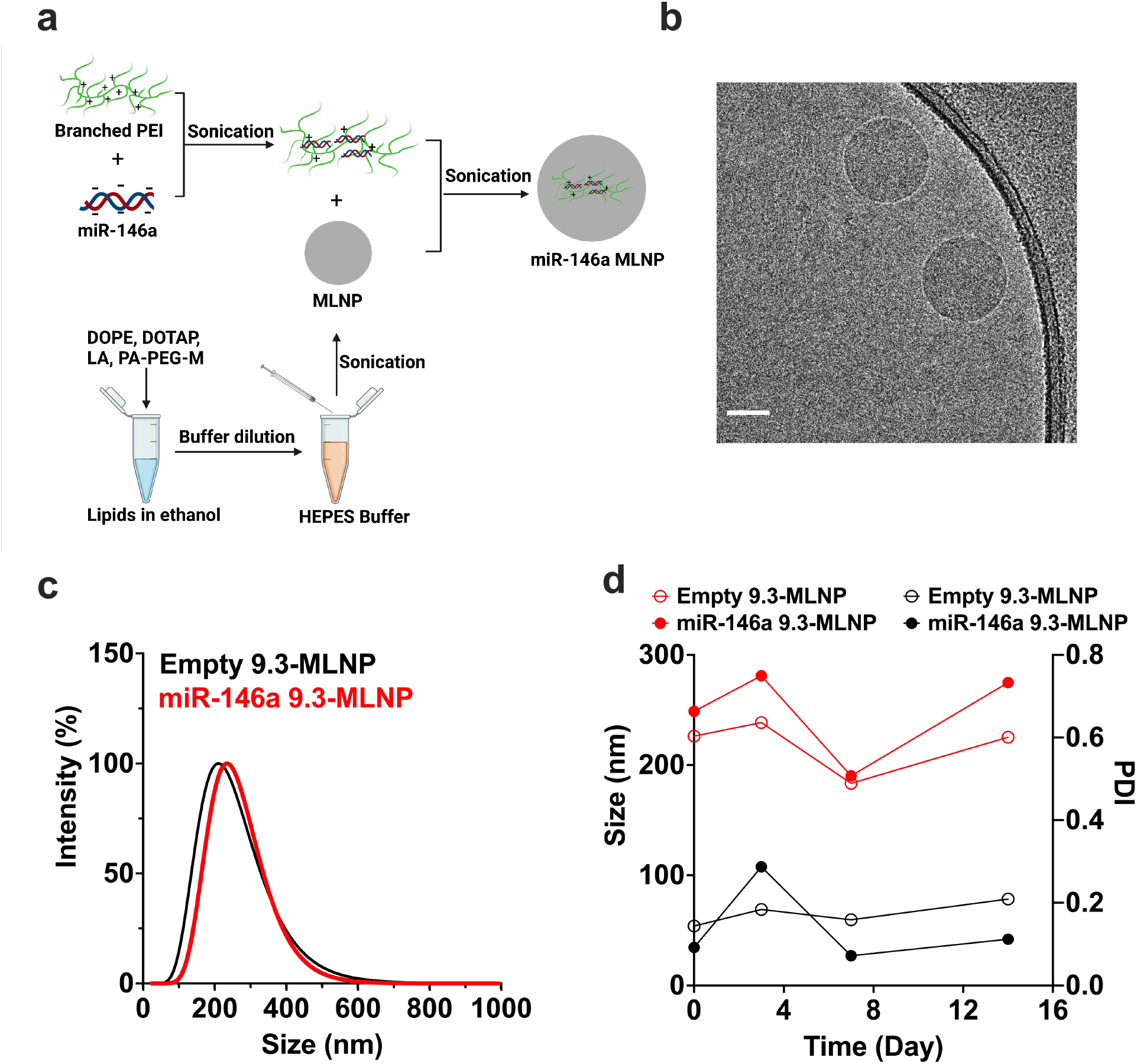
Fabrication and characterization of mannosylated lipid nanoparticle (MLNP). **a)** Schematic diagram of MLNP fabrication. Created with BioRender.com. **b)** Representative cryo-TEM image of miR-146a loaded MLNPs. Scale bar: 100nm. The experiment was performed twice. **c)** Representative size distribution of empty MLNPs and miR-146a loaded MLNPs. The experiment was performed three times. **d)** Storage stability in terms of size/diameter (left y-axis, red line) and polydispersity index (PDI) (right y-axis, black line) of empty MLNPs and miR-146a loaded MLNPs determined by dynamic light scattering (DLS) at time intervals of 0, 3,7, and 14 days.

**Table 1.** Physicochemical properties of mannosylated lipid nanoparticles.

| Formulations<sup>1</sup> | Size (nm)<sup>2</sup> | PDI<sup>3</sup> | Zeta potential (mV) |
| --- | --- | --- | --- |
| Empty 2-MLNP | 146.9 ± 3.5 | 0.158 ± 0.034 | -37.2 ± 7.0 |
| Empty 4.85-MLNP | 181.4 ± 23.8 | 0.138 ± 0.013 | -45.3 ± 4.7 |
| Empty 9.3-MLNP | 189.9 ± 13.5 | 0.140 ± 0.020 | -42.5 ± 1.5 |
| miR-146a 2-MLNP | 179.6 ± 5.4 | 0.212 ± 0.002 | 16.2 ± 5.9 |
| miR-146a 4.85-MLNP | 208.8 ± 2.3 | 0.137 ± 0.018 | 23.9 ± 3.9 |
| miR-146a 9.3-MLNP | 218.7 ± 13.2 | 0.181 ± 0.033 | 24.2 ± 4.7 |

<sup>1</sup> Formulation 2-MLNP, 4.85-MLNP, and 9.3-MLNP contains 2%, 4.85%, and 9.3% of mannose-conjugated (PA-PEG3-M) lipid respectively. <sup>2</sup> Size is expressed as the average diameter of particle size distribution by intensity. <sup>3</sup> PDI is the polydispersity index. The experiment was performed four times, except measurements for miR-146a 2-MLNP and miR-146a 4.85-MLNP were performed twice. Data are presented as mean ± SEM.

The entrapment efficiency (%EE) of miR-146a was assessed by agarose gel electrophoresis. The miR-146a loaded MLNPs without SDS treatment showed no visible band on the gel, while a clear band comparable to the size and intensity of the free miR-146a was observed after dissolving the miR-146a loaded MLNPs with SDS to release the entrapped miR, indicating a near 100% entrapment efficiency (**Figure S2a-b**). The encapsulation efficiency of fresh prepared (day 0) nanoparticle was further confirmed by Qubit™ microRNA assay that showed 100% EE (**Figure S2b**) which was stable over a period of 8 days at 4°C (**Figure S2b**), indicating that the miR-146a loaded MLNP was stable for 8 days when stored at 4°C.

### Mannosylated lipid nanoparticles efficiently deliver miR-146a to human alveolar macrophages

To evaluate the delivery of miR-146a to AMs using MLNPs, we measured the intracellular levels of mature miR-146a in primary human AMs after a 4 h treatment with untargeted LNPs loaded with scramble control, miR-146a loaded untargeted LNPs or miR-146a loaded mannosylated LNPs. We used a 4-fold lower miR-146a concentration (50 nM) compared to prior studies (200 nM)^6^. AMs treated with miR-146a loaded nanoparticles showed higher miR-146a levels compared to AMs treated with scramble-loaded nanoparticles (**Figure 2a**). The miR-146a levels in AMs delivered by MLNPs containing 9.3% mannosylated PA-PEG (9.3-MLNP) was significantly higher than those treated with MLNPs containing a lower amount of mannose-conjugated PA-PEG or those treated with untargeted LNPs (**Figure 2a**, **Figure S3a**). We also found the increase in miR-146a level was dose-dependent (**Figure 2b**). These results were repeated in additional donors as shown in **Figures S3a** and **S3b**. Prior studies have demonstrated that a minimum 100-fold increase in miR-146a is required to inhibit of pressure-induced inflammation in AMs *in vitro^6^*, therefore we selected the 50 nM concentration for further study (**Figure 2b**). To investigate whether the uptake of MLNPs was mannose receptor-dependent, we used mannose as a competitive inhibitor (**Figure 2c**). In the presence of excess mannose (20 mM), the uptake of 9.3-MLNP was reduced and comparable to that seen with untargeted LNPs.

**Figure 2.**
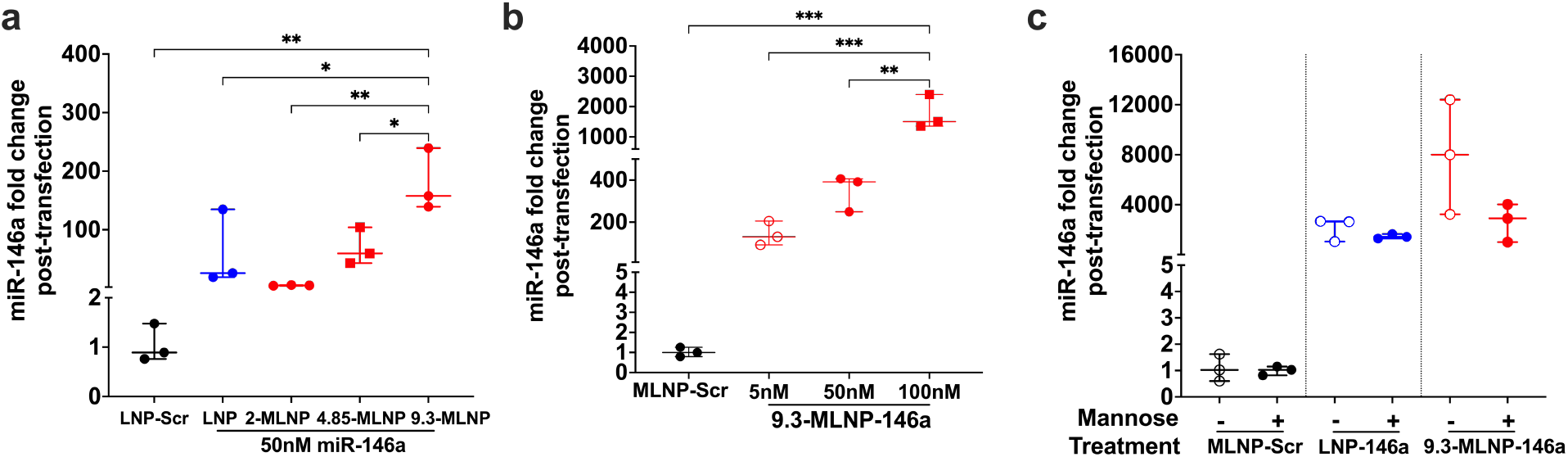
Cellular uptake of miR-146a loaded MLNPs in primary alveolar macrophages. **a)** Transfection efficiency in terms of miR-146a level in primary alveolar macrophages (AMs) obtained from donor 7 following delivery of miR-146a with different lipid nanoparticle formulations LNP, 2-MLNP, 4.85-MLNP and 9.3-MLNP (represented non-mannosylated LNP, mannosylated LNPs containing 2%, 4.85% or 9.3% of mannose-conjugated PA-PEG lipid respectively), calculated by △△Ct method, normalized to scramble-loaded LNP controls. Data are normally distributed by Shapiro-Wilk test. Statistical analysis was performed via one-way ANOVA with Tukey’s multiple comparisons test. **b)** Dose-dependent miR-146a level in AMs following 9.3-MLNP delivery of 5nM, 50nM or 100nM miR-146a, compared to scr-loaded 9.3-MLNP. Data are normally distributed by Shapiro-Wilk test. Data were analyzed by one-way ANOVA with Tukey’s multiple comparisons test. **c)** miR-146a level in AMs following delivery of miR-146a using LNPs or 9.3-MLNPs in the presence or absence of 20 mM mannose. Data were analyzed by Kruskal-Wallis with Dunn’s multiple comparisons test. Statistical differences are denoted as *p<0.05, **p<0.005, ***p<0.001. Data are presented as Min to Max. Show all points, n=3 wells per group. The experiment was performed twice.

### MLNP did not induce toxicity *in vitro* or *in vivo*

The cytotoxicity of 9.3-MLNPs was evaluated *in vitro* by MTS assay using primary human AMs and primary alveolar epithelial cells (i.e., pneumocytes). Cells were treated with empty 9.3-MLNPs at various nanoparticle concentrations (0 – 60 μg/mL) for 24 hours and the viability of both AMs and pneumocytes was unchanged in the range at nanoparticle concentrations up to 40 μg/mL (**Figure 3a-b**). These data indicate that the mannosylated LNPs were not cytotoxic at the concentrations (16 μg/mL *in vitro*, 15 μg per 25 g mouse) used in our studies.

**Figure 3.**
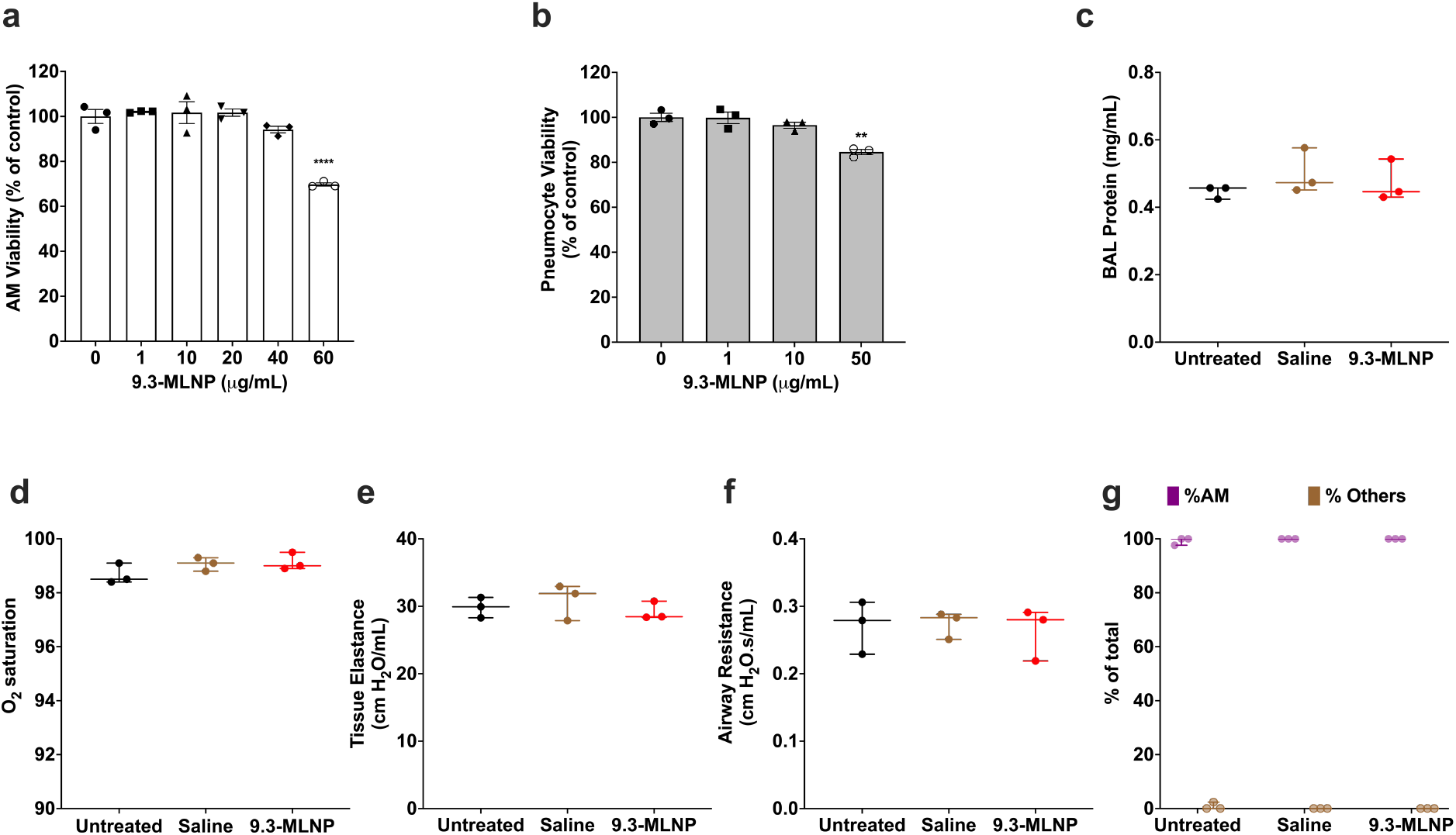
MLNPs do not induce toxicity on cells *in vitro* and on mice *in vivo*. Cell viability of **a)** AMs and **b)** pneumocytes after 24 h exposure to empty MLNP with various lipid concentrations by MTS assay. Cells with media alone were used as controls. Data were analyzed by one-way ANOVA with Tukey’s multiple comparisons test. **p < 0.005, ****p < 0.0001. Data are shown as means ± SEM, n=3 wells per group. **c)** The amount of total protein in bronchoalveolar lavage (BAL) fluid at 2 h after pulmonary administration of empty 9.3-MLNPs, untreated and saline-treated C57BL/6 mice were used as controls, n=3 per group. Data are not significant via one-way ANOVA with Tukey’s multiple comparisons test. Data are presented as Min to Max. Show all points. Effects of treatment of mice with saline or empty 9.3-MLNPs compared to untreated controls (n=3 per group) on **d)** blood oxygen saturation measured by pulse oximetry 2 h post-administration at initial (baseline) ventilation; **e)** lung tissue elastance; and **f)** airway resistance measurements at initial (baseline) ventilation. **g)** BAL differential cell counts in spontaneously breathing (SB) mice. Data are not significant via one-way ANOVA with Tukey’s multiple comparisons test. Data are presented as Min to Max. Show all points. Data are normally distributed by Shapiro-Wilk test.

To assess the *in vivo* toxicity of the 9.3-MLNPs, we administered empty 9.3-MLNPs to wildtype mice. Mice were sacrificed after 2 hours and the protein concentration in bronchoalveolar lavage (BAL) fluid was measured as a surrogate of lung barrier permeability. BAL protein levels were similar in mice treated with MLNPs compared to mice that received an intra-tracheal injection of saline or no treatment (**Figure 3c**). Lung function was also evaluated 2 h post-administration. Mice treated with MLNPs had normal oxygen saturations, lung tissue elastance, and airway resistance (**Figure 3d-f**). In addition, MLNPs did not induce pulmonary inflammation, as the majority of the cells from BALF were alveolar macrophages (**Figure 3g**). These data confirm that MLNPs do not induce toxicity *in vivo* within the short timeframe we tested.

### Mannosylated lipid nanoparticles preferentially deliver microRNAs to alveolar macrophages co-cultured with alveolar epithelial cells

To assess whether mannosylated lipid nanoparticles can be used to selectively deliver anti-inflammatory microRNAs to primary human lung macrophages, we developed a co-culture system where pneumocytes were grown with primary human AMs at an air-liquid interface. Co-cultures were treated with miR-scramble loaded untargeted LNP, miR-146a loaded untargeted LNP, miR-scramble loaded targeted MLNPs, or miR-146a loaded targeted MLNPs for 4 hours. Treatments were then removed, and cells were gently washed with PBS. After an additional 20 hours of incubation, cells were trypsinized and sorted by flow cytometry (**Figure S4**). We measured miR-146a levels in both the pneumocytes and AMs. While miR-146a levels were similar in pneumocytes and AMs treated with untargeted LNPs, AMs from co-cultures treated with miR-loaded MLNP had significantly higher miR-146a levels compared to pneumocytes (**Figure 4a-c**). Although the fold change varied by donor this effect was consistent using AMs from multiple donors. These results demonstrate that MLNPs preferentially deliver miR-146a to lung macrophages when they are cocultured with epithelial cells.

**Figure 4.**
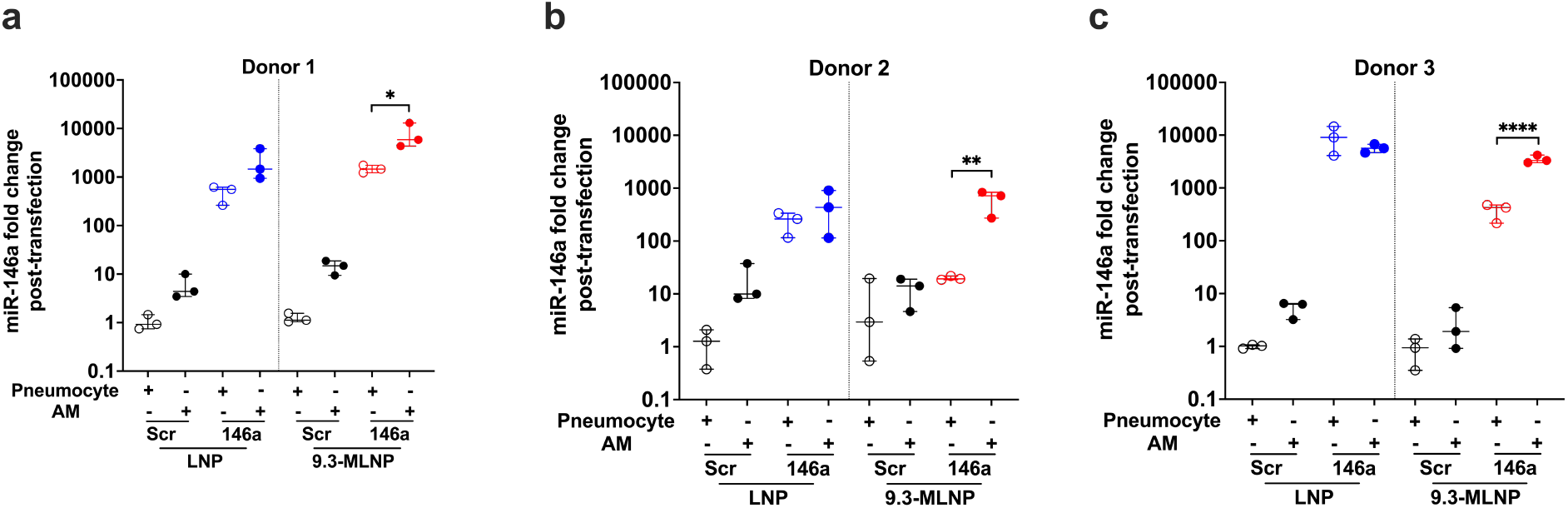
Mannosylated lipid nanoparticles preferentially deliver miR-146a to primary alveolar macrophages co-cultured with alveolar epithelial cells. miR-146a level following LNP or 9.3-MLNP delivery of miR-146a or scramble in flow-sorted pneumocytes and AMs from **a)** donor 1; **b)** donor 2; **c)** donor 3, n=3 wells per group. Data are normally distributed by Shapiro-Wilk test. Statistical analysis was performed separately on LNP data (left panel) and 9.3-MLNP data (right panel) by one-way ANOVA with Tukey’s multiple comparisons test. Statistical differences are denoted as *p<0.05, **p<0.01, ****p<0.0001. Data are presented as Min to Max. Show all points.

### Mannosylated lipid nanoparticle delivery of miR-146a potently dampens pressure-induced inflammation in an *in vitro* model of ventilator-induced lung injury

To characterize the inflammatory profile of AMs and pneumocytes in our co-culture system, we used a human cytokine array and measured cytokine levels in supernatants from pneumocytes and AMs subjected to an *in vitro* model of ventilator-induced lung injury from increased transmural pressure (i.e., Barotrauma). We found that IL8, MIP-1a, and MIP-1ß were increased in AM supernatants but not those from pneumocytes (**Figures S5a-b**). In contrast, IL6 was secreted by pneumocytes in response to barotrauma to a much greater degree than in AMs.

To evaluate the efficacy of miR-146a loaded MLNPs to diminish pro-inflammatory cytokines/chemokines secretion compared to untargeted LNPs, we measured the production of MIP-1a, IL-8 and IL-6 in response to barotrauma in our co-culture system. Scramble nanoparticle-treated co-cultured cells exhibited increased levels of MIP-1a, IL-8, and IL-6 (left panel of **Figure 5a-c**) when exposed to *in vitro* barotrauma compared to control cells not exposed to pressure. Following treatment with miR-146a-loaded LNPs or MLNPs, co-cultured cells had decreased cytokine secretion. Specifically, treatment with miR-146a-loaded MLNPs led to significantly less production of the AM-derived cytokines, MIP-1a and IL-8 (right panel **Figure 5a-b**) compared to cells treated with untargeted LNPs. Cells treated with either untargeted LNPs or MLNPs also had decreased levels of the epithelial-derived cytokine IL-6 (right panel of **Figure 5c**) when subjected to *in vitro* barotrauma; however, miR-146a-loaded LNPs and MLNPs decreased this epithelial-derived cytokine to a similar extent. These data demonstrate that overexpression of miR-146a by MLNPs can more potently mitigate pressure-induced MIP-1a and IL-8 secretion by AMs in co-culture compared to untargeted nanoparticles. This ability to specifically dampen the production of AM-derived cytokines in co-cultured cells strongly suggests that MLNPs preferentially target lung macrophages.

**Figure 5.**
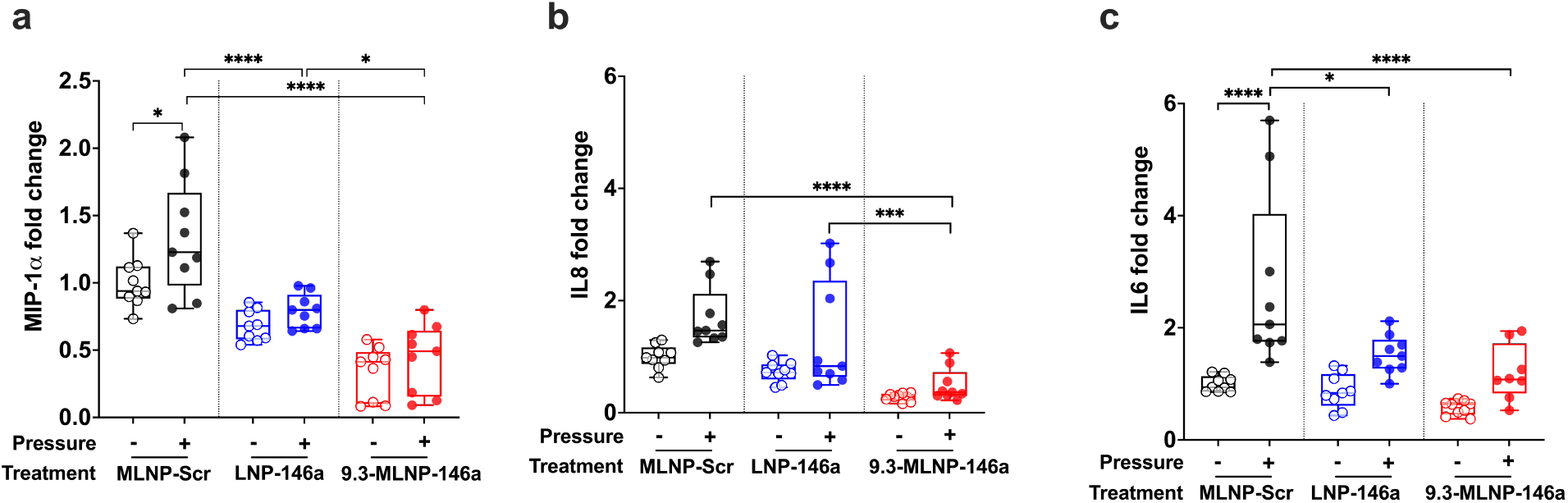
MLNP delivery of miR-146a more potently dampens pressure-induced inflammation in co-culture. MIP-1 alpha (MIP-1 a), IL8, and IL6 levels were determined following treatment with miR-146a loaded LNPs or 9.3-MLNPs in co-culture of pneumocytes and alveolar macrophages (AMs) from three donor lungs (donor 3, 4, 5) subjected to 16 h of oscillatory pressure at the air-liquid interface, normalized to unpressurized scramble-loaded MLNP treated controls. n = 3 for each donor. **a)** Fold change in (MIP-1a) secretion; Data are normally distributed by Shapiro-Wilk test. Data were analyzed via one-way ANOVA with Sidak’s multiple comparisons test. **b)** Fold change in IL8 secretion; Data are lognormally distributed by Shapiro-Wilk test. Data were analyzed via one-way ANOVA with Sidak’s multiple comparisons test. **c)** Fold change in IL6 secretion; Data are lognormally distributed by Shapiro-Wilk test. Data were analyzed via one-way ANOVA with Sidak’s multiple comparisons test. Statistical differences are denoted as *p<0.05, **p<0.005, ***p<0.0005, ****p<0.0001. Data are presented as Min to Max. Show all points.

### *In vivo* delivery of mannosylated lipid nanoparticles to lung macrophages

To assess the ability of MLNPs to target lung macrophages *in vivo*, wild-type mice were administered Cy3-labeled scramble-miR loaded LNPs or MLNPs by intra-tracheal injection. We have previously demonstrated that this technique delivers nanoparticles to the distal lung^6^. Four hours after nanoparticle delivery, cryo-sectioned lung tissues were stained with a macrophage marker (CD68) (**Figure 6a, b**). The degree of CD68 and Cy3 co-localization was assessed by Image J and showed that mice treated with Cy3-scr loaded MLNPs had increased macrophage area that contained Cy3-scr compared to those treated with untargeted LNPs (**Figure 6c**). To increase the rigor of this finding, we also determined the co-occurrence by manually counting the number of macrophages containing nanoparticles (**Figure 6d**) which was performed by another investigator who was blinded to the treatment groups. These data demonstrate that MLNPs preferentially distribute into lung macrophages in mice.

**Figure 6.**
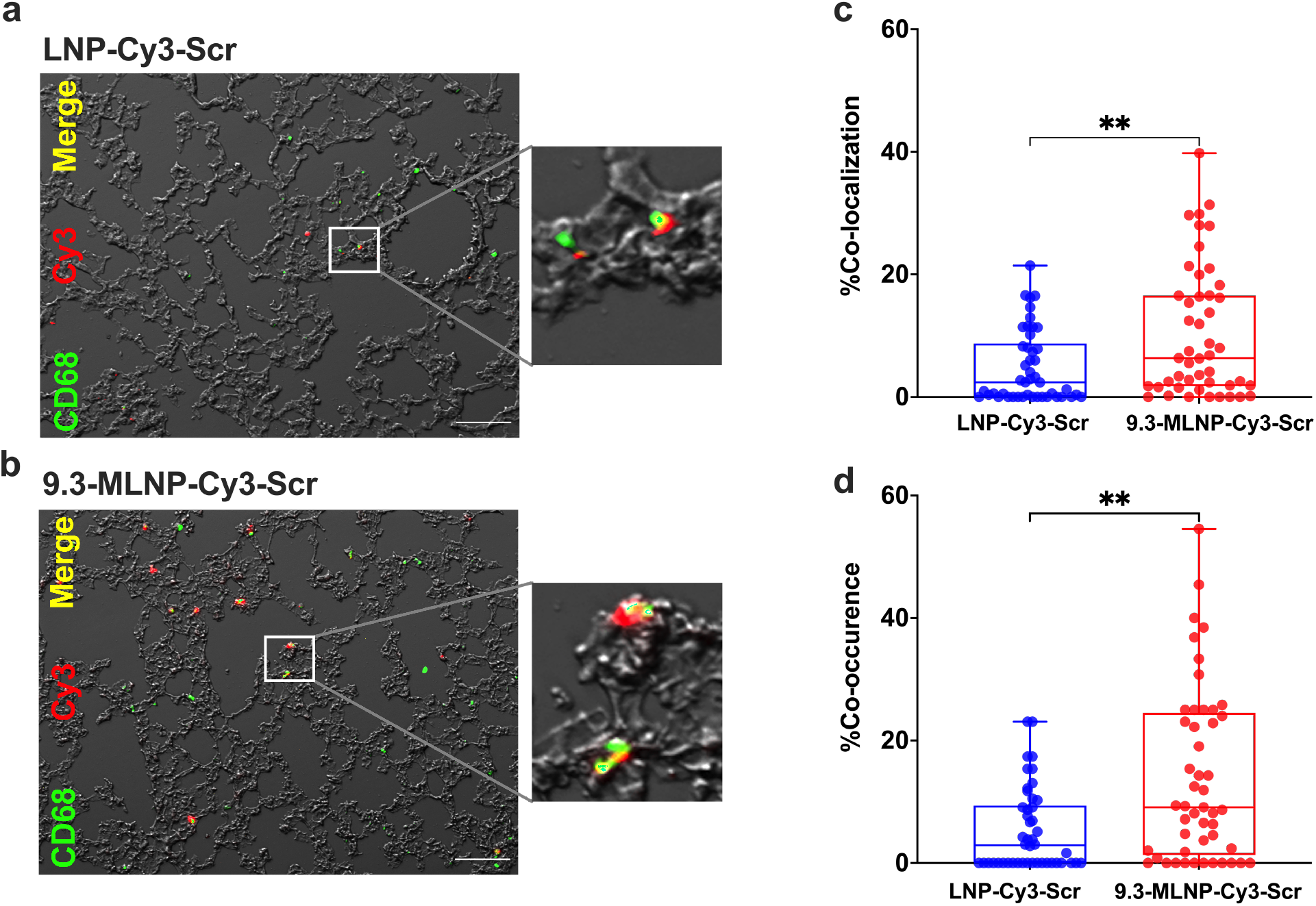
In vivo lung cell-distribution of Cy3-Scramble loaded MLNP. **a)** Representative fluorescence and DIC images from lung tissue of spontaneously breathing mice treated with Cy3-scramble-miR loaded LNPs or **b)** Cy3-scramble-miR loaded 9.3-MLNPs by immunofluorescence staining for CD68 (green). Scale bar: 100 μm, n=3 mice per group. **c)** Percentage macrophage cell area containing Cy3-scramble loaded nanoparticles using immunofluorescence images. n=19 high-power fields from one animal per group and n=15 high-power fields from the other two animals per group. **p=0.0055 by Mann-Whitney test. Data are presented as Min to Max. Show all points. **d)** Percentage of the number of macrophages containing Cy3-scramble loaded nanoparticles. n=19 high-power fields from one animal per group and n=15 high-power fields from the other two animals per group. **p=0.0013 by Mann-Whitney test. Data are presented as Min to Max. Show all points.

### Mannosylated lipid nanoparticle delivery of miR-146a mitigates lung injury in a model of hemorrhagic shock-induced acute respiratory distress syndrome

To test the efficacy of MLNPs to prevent lung inflammation, we developed a model of acute respiratory distress syndrome (ARDS) due to the combination of hemorrhagic shock and mechanical ventilation. ARDS from hemorrhagic shock (HS-ARDS) is common in civilian and military populations and there are no targeted therapies to treat excessive inflammation in these patients. Wild-type mice were subjected to 30 minutes of hemorrhagic shock by removing blood from an arterial catheter until the mean arterial pressure reached 30-40 mm Hg. Mice were then resuscitated with the shed blood and isotonic fluid and placed on mechanical ventilation for 2 hours. HS-ARDS led to significantly higher levels of IL-6 and CXCL1/KC (mouse analog of IL-8) in bronchoalveolar lavage (BAL) fluid compared to animals exposed to HS alone (**Figures S6a-b**). This indicates that while HS alone may not cause excessive inflammation, the physical forces generated during mechanical ventilation are the primary driver of inflammation in this model. Mice subjected to HS-ARDS had increased lung elastance (i.e., stiffness, **Figure S6c**) and significantly lower oxygenation saturations (**Figure S6d**) compared to mechanically ventilated mice that did not undergo HS. miR-146a levels in BAL cells from HS-ARDS mice were similar to those found in controls despite the increase in inflammatory cytokines (**Figure S6e**). miR-146a has a role in dampening lung inflammation; however, prior work has demonstrated that supraphysiologic increases to > 100-fold are required to mitigate lung injury/inflammation. The failure to upregulate miR-146a level in HS-ARDS mice suggests that a strategy to increase miR-146a in lung macrophages may have a therapeutic benefit.

To test this hypothesis wild-type mice were treated with untargeted miR-146a loaded LNPs, targeted miR-146a loaded MLNPs, or scramble-loaded control MLNPs *via* intratracheal administration after hemorrhage shock but just prior to mechanical ventilation. Prior to these *in vivo* studies, a dose-response study was performed by measuring the miR-146a levels 2 hours following administration of 0.1 nmol or 1 nmol miR-146a loaded MLNPs. We found mice treated with 0.1 nmol miR-146a loaded MLNPs had nearly a 10,000-fold increase in miR-146a levels in BAL cells (**Figure S7**). Since previous studies indicate that this level of miR-146a over-expression was sufficient to dampen lung injury and inflammation *in vivo^6^*, all mice were treated with 0.1 nmol of miR-146a loaded nanoparticles. AMs recovered from BAL following HS had a robust increase in miR-146a levels with both targeted and untargeted LNPs (**Figure 7a**). To demonstrate the functionality of the delivered miR-146a we measured levels of a well-known target transcript of this miR (i.e., TRAF6) and found that both targeted and untargeted LNPs significantly reduced TRAF6 levels compared to mice treated with scramble control (**Figure 7b**). The miR-146a level in whole lung homogenate increased about 20-100-fold (**Figure S8**) which was much lower compared to the increases in miR-146a in BAL AMs. Following HS-ARDS, mice treated with miR-146a loaded MLNPs had significantly less BAL IL-6 as compared to mice treated with scramble-loaded MLNPs or mice treated with miR-146a loaded LNPs. In addition, mice treated with miR-146a loaded MLNPs exhibited significantly less CXCL1/KC levels as compared to mice treated with scramble-loaded MLNPs (**Figure 7c, d**). BAL protein levels were measured as a marker of lung barrier permeability and were not affected by miR-146a treatment at this time point compared to scramble controls (**Figure 7e**). Although lung stiffness was not significantly changed in miR-146a nanoparticle-treated mice compared to scramble-nanoparticle control-treated mice, the change in lung elastance of miR-146a-MLNP treated mice was lower compared to that in miR-146a-LNP treated mice (**Figure 7f**). There were no significant differences in oxygenation or in BAL inflammatory cell counts in miR-146a-MLNP treated mice (**Figure 7g-h**). These data indicate that delivery of miR-146a to AMs by MLNPs can dampen inflammation in a clinically relevant model of HS-ARDS.

**Figure 7.**
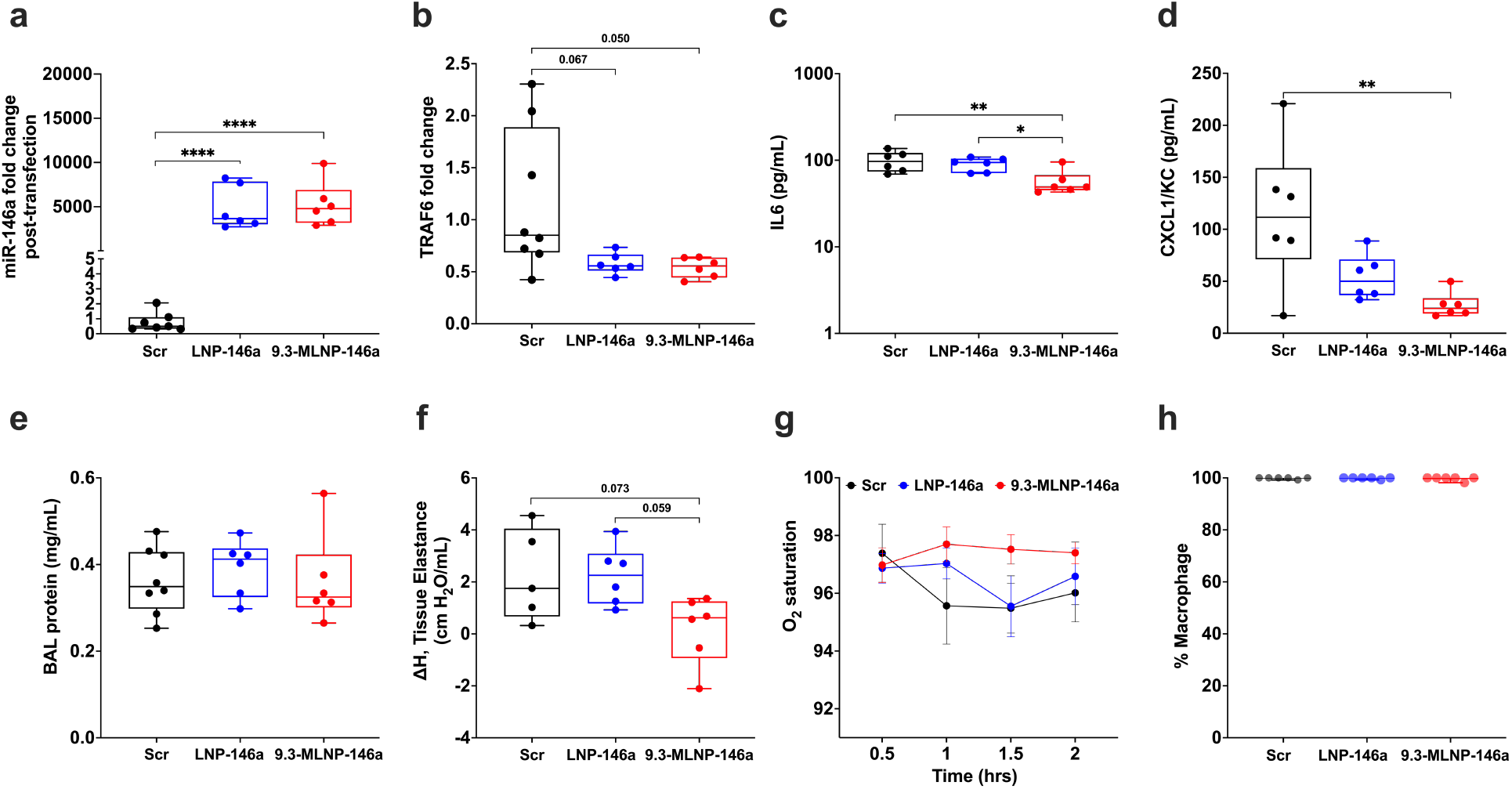
MLNP delivery of miR-146a mitigates lung injury in a model of hemorrhagic shock-induced ARDS. **a)** miR-146a level in RNA from BAL cell pellet following nanoparticle delivery in HS-ARDS mice. Relative expression determined by ΔΔCt method, normalized to scramble control. Data are lognormally distributed by Shapiro-Wilk test. ****p<0.0001 via oneway ANOVA with Tukey’s multiple comparison test. **b)** TRAF6 mRNA levels from BAL cells by ΔΔCt method normalized to scramble. Data are normally distributed by Shapiro-Wilk test. Data were analyzed by one-way ANOVA with Tukey’s multiple comparison test. **c)** BAL IL6 level from miR-146a or scramble (scr) loaded nanoparticles treated HS-ARDS mice. Data are lognormally distributed, *p<0.05, **p<0.005 via one-way ANOVA with Tukey’s multiple comparison test. and **d)** BAL IL8 level from miR-146a or scramble (scr) loaded nanoparticles treated HS-ARDS mice. Data are normally distributed by Shapiro-Wilk test. **p<0.01 via oneway ANOVA with Tukey’s multiple comparison test. **e)** BAL protein concentration from miR-146a and scr nanoparticles treated HS-ARDS mice. Data are normally distributed by Shapiro-Wilk test. No significance via one-way ANOVA with Tukey’s multiple comparison test. **f)** Change in lung elastance during 2 h period of ventilation following treatment with miR-146a or scr loaded nanoparticles. Data are normally distributed by Shapiro-Wilk test. Data were analyzed via one-way ANOVA with Tukey’s multiple comparison test. **g)** Oxygenation throughout duration of ventilation measured via pulse oximetry. Data are normally distributed by Shapiro-Wilk test. Data were analyzed via repeated measures two-way ANOVA with Tukey’s multiple comparison test. **h)** Percentage macrophages in BAL cells by differential counts after ventilation. For all panels, n=8 for scramble group and n=6 for miR groups. Data are presented as Min to Max. Show all points.

## Discussion

The goal of this study was to design and fabricate lipid nanoparticles that specifically target lung macrophages to facilitate the delivery of anti-inflammatory microRNAs to these cells and thereby more effectively mitigate ventilation induced lung injury during the acute respiratory distress syndrome. To achieve this goal, we synthesized nanoparticles with mannosylated lipids (MLNP) at the surface to preferentially deliver miR-146a to alveolar macrophages (AMs). We demonstrated that these MLNPs robustly increase miR-146a levels in human primary AMs *in vitro* in a mannose-dependent fashion (**Figure 2**). We then used a co-culture system that models the lung microenvironment and showed that MLNPs were preferentially delivered to AMs (**Figure 4**) and decreased the production of macrophage-specific cytokines/chemokines (**Figure 5**). MLNPs were then delivered to mice *via* intra-tracheal injection. MLNPs were preferentially delivered to AMs in mouse lungs and no evidence of toxicity was observed (**Figure 3** & **Figure 6**). Finally, in a proof-of-principle experiment, we demonstrated that pulmonary delivery of miR-146a loaded MLNPs dramatically increased miR-146a levels in AMs, decreased the expression of a well-described target (ie. TRAF6), and decreased lung inflammation in a clinically relevant model of ARDS from mechanical ventilation in the setting of hemorrhagic shock (**Figure 7**).

Other groups have used a similar strategy to deliver nanoparticles to lung macrophages. A study conducted by Wijagkanalan et al. demonstrated that intra-tracheal administration of mannosylated liposomes improved targeting efficiency to alveolar macrophages in rats ^15^. A more recent study showed mannose-functionalized solid lipid nanoparticles loaded with the tuberculosis antibiotic isoniazid had higher accumulation in murine alveolar macrophages^21^. Other ligands used for AM targeting include maleylated bovine serum albumin (MBSA) and *O*-steroyl amylopectin (*O*-SAP)^9^. Liposomes coated with MBSA, a ligand specific for alveolar macrophage scavenger receptors, preferentially accumulate in the alveolar macrophages after aerosol delivery^22^. *O*-SAP-modified nanoparticles also showed improved delivery to alveolar macrophages^23^. Ultimately, we decided to use mannose functionalized nanoparticles due to their low immunogenicity^17^ and low toxicity^24^. Despite the continued development of nanoparticle-based drug delivery systems, most of the AM-targeting studies are still in the early pre-clinical stage and have been used primarily to deliver antibiotic agents^25, 26^. To our knowledge, macrophage-targeted nanoparticles have not been used previously to deliver anti-inflammatory microRNAs that can mitigate lung injury in a clinically relevant model of ARDS.

One of the potential advantages of targeted drug delivery to a specific cell type is the ability to reduce dose-limited toxicities, the initial drug dose, and dosing frequency by increasing local concentrations in the lung^7, 27, 28^. Our targeted MLNP delivery of 50 nM miR-146a to AMs *in vitro* resulted in greater inhibition of pressure-induced proinflammatory cytokine release compared to untargeted LNP delivery (**Figure 2b** & **Figure *5***). This miR-146a concentration (50 nM) was significantly lower than the dose required in our prior work (200 nM) to reduce *in vitro* proinflammatory cytokine release using untargeted LNPs^6^. Interestingly, the miR-146a levels in BAL AMs following delivery by either targeted MLNPs or untargeted LNP *in vivo* were similar (**Figure 7a**). This may indicate that *in vivo*, following injury from HS-ARDS, AMs exhibit similar uptake of targeted and untargeted nanoparticles, and the *in vivo* conditions under which MLNPs are preferentially taken up by AMs require further investigation. Despite similar miR-146a levels in mice treated with both targeted and untargeted LNPs, mice treated with targeted MLNPs had decreased lung inflammation and improved lung function compared to mice treated with untargeted LNPs. A single dose of only 0.1 nmol miR-146a targeted MLNPs immediately prior to starting mechanical ventilation was required to achieve this effect. This was significantly lower than the previously reported regimen of two doses of 1 nmol miR-146a using untargeted LNPs to mitigate lung inflammation from mechanical ventilation *in vivo*. Furthermore, the first dose in this prior study was given 24 hours prior to the initiation of mechanical ventilation which is often not feasible in patients with ARDS. The ability to use a lower dose of miR and administer at the time mechanical ventilation is an important advance in this field and increases the translational potential of targeted MLNP delivery of microRNA therapeutics to mitigate ventilation-induced lung injury during ARDS.

Although our targeted MLNPs enhanced delivery to alveolar macrophages by targeting the mannose receptors on the surface of AMs, the mannose receptor is also expressed on hepatic endothelial cells and dendritic cells (DCs)^29^. Our study used the pulmonary delivery route which theoretically should bypass the liver; however, there is a possibility that these nanoparticles could enter the systemic circulation and have off-target effects due to mannose receptor expression in cells of other organs. Although DCs make up a relatively small proportion of lung immune cells, prior studies have demonstrated that they play a key role in lung injury^30–33^. During hemorrhagic shock^34^- or LPS-induced ARDS^35^, the number of DCs in lung tissue is increased. There are also some studies that suggest miR-146a is a regulator of inflammation in microvascular endothelial cells^36^ and we have not specifically studied this cell type. Therefore, future studies should investigate how targeting the mannose receptor to deliver miR-146a may affect different cell types. Our current study focused on miR-146a delivery to the distal alveolus, but the alveolar-capillary barrier becomes permeable during ARDS. This leads to hetergeous occlusion of injured alveoli and may hinder the transport of nanoparticles from the upper airways to the site of injury. Therefore, further research is needed to determine how to enhance nanoparticle delivery to these occluded lung regions. Another limitation of our current study is that nanoparticles were administered *in vivo* via intratracheal injection which has limited clinical applicability. However, previous studies have demonstrated that nanoparticles can be delivered by aerosolization^37–40^. In fact a liposomal inhalation formulation of an antibiotic was recently approved by FDA^41^ for patients with certain types of infection. Therefore, future studies could investigate whether MLNPs can be formulated for delivery using nebulization.

In summary, we have developed targeted nanoparticles (MLNPs) that showed enhanced delivery of miR146a to lung macrophages. Delivery of this microRNA to lung macrophages ameliorated ventilation-induced lung injury during hemorrhagic shock-induced ARDS. Importantly, we were able to achieve supraphysiological miR-146a expression (10,000-fold) using a single 0.1 nmol dose just prior to mechanical ventilation. This nanoparticle system with mannose conjugation holds the potential to achieve therapeutic effects with a single dose at the time mechanical ventilation is initiated. In addition, macrophage-targeted delivery of miR146a allows rapid onset of action leading to rapid clinical response which is critical for early prevention or treatment of lung injury during ARDS. Future studies will be needed to explore the therapeutic potential of miR-146a MLNPs to mitigate ventilator-induced lung injury in patients with ARDS.

## Materials and methods

### Materials

1,2-Dioleoyl-sn-glycero-3-phosphoethanolamine (DOPE), 1,2-dioleoyl-3-trimethylammonium-propane (DOTAP), 1,2-dipalmitoyl-sn-glycero-3-phospho((ethyl-1’,2’,3’-triazole)triethyleneglycolmannose) (ammonium salt) (PA-PEG3-mannose) were purchased from Avanti Polar Lipids (Alabaster, AL, USA). Linoleic acid (LA), D-α-Tocopherol polyethylene glycol 1000 succinate (TPGS) and polyethylenimine (PEI, 2,000 MW) were obtained from Sigma Aldrich (Natick, MA, USA). D-(+)-mannose was also purchased from Sigma Aldrich. RPMI-1640 medium was purchased from Sigma Aldrich, Gibco Opti-MEM reduced serum medium was purchased from ThermoFisher (Waltham, MA, USA), human epithelial cell (HEC) media with the supplemental kit was purchased from Cell Biologics Inc. (Chicago, IL, USA).

### Animal use

All animal studies were approved by the Institutional Animal Care and Use Committee (IACUC) at The Ohio State University under protocols 020A00000037 and 2011A00000081-R3. Male wild-type C57Bl/6J mice were purchased from the Jackson Laboratory (Bar Harbor, ME, USA). All animal experiments were carried out in compliance with all guidelines and ethical regulations.

### Fabrication of mannose-conjugated lipid nanoparticles (MLNPs)

MLNPs were fabricated by the ethanol injection and sonication method as described previously^6^. Briefly, a mixture of lipids including DOPE, DOTAP, LA, and PA-PEG3-mannose were dissolved in ethanol at a molar ratio of 40:10:48:2 or 40:10:48:5 or 40:10:48:10. The mixture was injected into 20mM HEPES buffer and then sonicated to form empty MLNPs containing 2%, 4.85% and 9.3% PA-PEG-mannose respectively. Empty LNPs were prepared using the same method with a molar ratio of 40:10:48:2 for DOPE:DOTAP:LA:TPGS. miR-146a or miR scramble control (ThermoFisher Scientific, Waltham, MA) were mixed with PEI at an N:P ratio of 25 (the ratio of moles of PEI amine groups to nucleic acid phosphate groups) in HEPES buffer and incubated at room temperature for 10 min to form PEI/miRNA polyplexes. Two different miR-146a mimics (Ambion/Thermo Pre-miR™ and miRVana mimic) were used. The empty nanoparticles and polyplex solutions were then mixed at a lipid-to-nucleic acid final mass ratio of 10 and this mixture was sonicated for 5 min and then incubated for 10 min at room temperature. To perform the *in vivo* distribution study, a Cy3-Dye-labeled miR scramble control (ThermoFisher Scientific) was loaded into the MLNPs using the same method as described above.

### Physico-chemical characterization of MLNP

Particle size and polydispersity index (PDI) were measured *via* dynamic light scattering (DLS) using a NICOMP Z3000 Nano DLS/ZLS Systems (Entegris, Brillerica, MA, USA). Nanoparticles were diluted to a ratio of 1:100 with nuclease-free water. Zeta potential was determined by laser Doppler velocimetry (LDV) using NICOMP Z3000 Nano DLS/ZLS Systems. Experiments were performed four times.

### Encapsulation efficiency

Encapsulation efficiency was determined by agarose gel electrophoresis as described previously^6, 42, 43^. Briefly, miR-146a-loaded MLNPs were lysed with 0.5% sodium dodecyl sulfate (SDS) (BioRad, Hercules, CA), and loaded onto 2% agarose (Fisher BioReagents) gel containing Labsafe nucleic acid stain (GBiosciences). miR-146a-loaded MLNPs without SDS were also loaded into the gel for comparison. Gel electrophoresis was run at 100 V for 45–60 min. Images of the gel were captured under UV light using ChemiDoc Imaging System (Bio-Rad). Encapsulation efficiency was quantified by Qubit™ microRNA Assay Kits (ThermoFisher Scientific) according to the manufacturer’s protocol. Briefly, a standard curve was generated using the microRNA standards from the kit. miR-146a loaded MLNPs or empty MLNPs were lysed using 1% TritonX-100 (Sigma) and diluted in a working solution. Intact nanoparticles without adding 1% TritonX-100 were also diluted in a working solution. Following vigorous vortexing, samples were incubated at room temperature for 2 min to reach optimal fluorescence, protected from light. 200 μL of samples were transferred to a 96-well plate, and fluorescence intensity was read at excitation 500 nm and emission 520 nm. The final TritonX-100 (Sigma) in the assay was 0.001% (v/v) as recommended by the manufacturer’s instruction. The concentration of miR-146a in MLNPs was determined using the equation generated from the standard curve. Encapsulation efficiency was calculated by dividing the absolute difference of miR-146a concentration between lysed nanoparticles and intact nanoparticles by lysed nanoparticles.

### Cryo-transmission electron microscope (Cryo-TEM)

The morphology of the miR-146a loaded MLNPs was assessed on a Thermo Scientific Glacios Cryo-TEM instrument. Briefly, Cryo-TEM samples were prepared by adding 3 μl of the miR-146a loaded MLNPs onto a 400-mesh copper lacey carbon grid. The grid was plunge-frozen in liquid ethane using the Vitrobot system. Images were acquired using Glacios Cryo-TEM.

### Physical stability of MLNPs

Empty MLNPs and miR-146a loaded MLNPs were stored at 4°C and particle size and PDI were measured at days 0, 3, 7, and 14. microRNA content in MLNPs over time was determined by Qubit™ microRNA Assay Kits (ThermoFisher Scientific) according to the manufacturer’s protocol.

### Cell culture

#### Monoculture

Human primary alveolar macrophages (AMs) were isolated *via* ex vivo lavage from de-identified donor lungs deemed unsuitable for transplantation obtained from Lifeline of Ohio Organ Procurements agency (Columbus, OH). All lungs were from subjects with no history of chronic lung disease or cancer and were non-smokers for at least 1 year. After the collection of bronchoalveolar lavage (BAL) fluid, BAL was centrifuged at 800 rcf for 10 min at 4°C. RBCs were lysed using a 1x RBC lysis buffer, the reaction was stopped by adding RPMI media supplemented with 10% Fetal Bovine Serum (FBS) and 1% antibiotic-antimycotic (anti-anti) (Thermo Fisher Scientific). Cells were centrifuged again at 800 rcf for 10 min at 4°C, and the cell pellet was resuspended and frozen down in FBS with 10% DMSO. Prior to experiments, AMs were rapidly thawed and cultured in RPMI media supplemented with 10% FBS and 1% anti-anti at 37°C in a humidified, 5% CO_2_ incubator. For transfection, cells were cultured in Opti-MEM (with 1% anti-anti). Primary human pneumocytes (Cell Biologics Inc, Chicago, IL, USA) were cultured in human epithelial cell (HEC) growth media supplemented with a human epithelial supplement kit.

#### Co-culture

Pneumocytes were seeded on Transwell 6-well 24mm inserts (Corning Inc., NY, USA) at a density of 1.5 x 10^5^ cells/insert in HEC growth media and incubated for 48 h at 37°C and 5% CO_2_. HEC growth media from the apical chamber was then removed and cells were washed with 1X Ca^2+^/Mg^2+^-free Dulbecco’s phosphate-buffer saline (PBS) (Gibco). AMs were concentrated into 10 μL Opti-MEM (with 1% anti-anti) and seeded on top of pneumocytes at a cell density ratio of 1:1^44^. AMs were allowed to adhere for 1 h. For the transfection experiment, HEC growth media from the basal chamber was replaced with FBS-free HEC basal media (with 1% anti-anti).

### *In vitro* cytotoxicity (MTS assay)

The cytotoxicity of MLNP was evaluated by MTS assay using CellTiter 96^®^ AQ_ueous_ reagents according to the manufacturer’s protocol (Promega, Madison, WI, USA) and as previously described^45^. Primary AMs were seeded into 96-well plates with clear flat-bottomed wells at a density of 1 x 10^4^ cells in 100 μL RPMI media supplemented with 10% FBS and 1% anti-anti per well. AMs were allowed to adhere for 1 h. AMs were then washed gently with warm DPBS and treated in triplicate with different concentrations of MLNPs for 24 h (humidified, 37 °C, 5% CO_2_). After 24 h incubation, the cell medium was aspirated and exchanged with 100 μl medium containing a final concentration of 333 μg/ml MTS and 25 μM PMS. AMs were incubated for 4 h at 37 °C. The absorbance was measured at 490 nm with a microplate reader.

### RNA extraction

RNA from cells was extracted using the standard phenol–chloroform RNA extraction and purification method by TRIzol (Ambion, Carlsbad, CA, USA) according to the manufacturer’s protocol. RNA from tissues was extracted and purified using Direct-Zol™ RNA MiniPrep Plus kit (Zymo Research, Irvine, CA, USA).

### Quantitative real-time PCR (RT-qPCR)

RT-qPCR with TaqMan primer/probes was used to determine relative expression levels of miR-146a compared to U18 (human) or sno251 (mouse) endogenous control genes (ThermoFisher, assay IDs: 478399_mir, 001204, 001236, respectively). TRAF6 relative expression was determined relative to GAPDH control (ThermoFisher, assay IDs: Mm00493836_m1 (TRAF6), Mm99999915_g1 (GHPDH)). Following RNA extraction and purification, cDNA was synthesized using a high-capacity cDNA reverse transcription kit (ThermoFisher) on Biored Thermocycler (PTC-200, Peltier Thermal Cycler). qPCR was then performed with TaqMan Universal PCR Master Mix (ThermoFisher) on QuantStudio 3 (ThermoFisher) and data were quantified by the 2^-ΔΔCT^ method.

### *In vivo* toxicity of MLNP

Wild-type C57Bl/6J mice were anesthetized and intratracheally administered with 15 μL sterile saline and 15 μL empty 9.3-MLNPs containing 15 μg lipids. Spontaneously breathing untreated mice were used as controls. 2 h after nanoparticle administration, mice were anesthetized and a tracheostomy cannula was placed, and mice were connected to a Flexivent small animal ventilator (Scireq, Montreal, QC, Canada). Baseline lung physiology measurements were obtained by performing a recruitment maneuver followed by forced oscillation to determine tissue elastance. Blood oxygenation was monitored using a mouse thigh pulse oximeter (Starr Life Sciences Corporation, Oakmont, PA, USA). Mice were then euthanized, and BAL was performed by instilling 1 ml of saline into the lungs twice and withdrawing it each time. BAL fluid was then centrifuged for 10 min at 2000 rpm at 4°C and the supernatant was collected for subsequent analysis. Red blood cells were lysed with RBC lysis buffer. BAL cells were counted with a TC20 Automated Cell Counter (Bio-Rad Laboratories, Hercules, CA, USA), and a differential stain was performed (Hema 3 Stat Pack, ThermoFisher Scientific) according to the manufacturer’s instruction. The remaining cells were combined and pelleted and then resuspended in TRIzol (Qiagen) for RNA extraction. Following BAL, the left lung was inflated and fixed with 10% formalin. Other organs including the right lungs, liver, heart, spleen, kidney, and diaphragm were harvested and snap-frozen in liquid nitrogen.

### *In vitro* uptake of MLNPs

Primary AMs were seeded on Transwell 6-well inserts at a density of ~1 x 10^6^ cells per insert and allowed to adhere for 1 h in complete RPMI media supplemented with 10% FBS and 1% anti-anti at 37°C in a humidified, 5% CO_2_ incubator. Following adherence, media was removed and replaced with Opti-MEM supplemented with 1% anti-anti in the basal chamber, and media was removed from the apical chamber and washed once gently with warm DPBS. AMs were treated with miR-146a loaded MLNPs and LNPs in Opti-MEM (with antibiotic supplement) at miR-146a concentration of 50 nM for 24 h. AMs treated with miR-scramble-loaded nanoparticles were used as a control. Cellular RNA was isolated for quantitative real-time (RT-qPCR) analysis.

### Mannose competition study

AMs were pre-treated with 20 mM mannose (stock dissolved in HEPES buffer) in RPMI media for 1 h^46^. An equal volume of HEPES buffer was added to wells not containing mannose. The mannose solution was removed after 1 h and AMs were gently washed twice with DPBS (Gibco). AMs were treated with miR-146a loaded MLNPs and LNPs in Opti-MEM (with 1% anti-anti) at miR-146a concentration of 50 nM for 24 h. For wells pretreated with 20 mM mannose, the same concentration of mannose was also included together with the nanoparticle treatments. Cellular RNA was isolated for RT-qPCR analysis. All treatments were performed in triplicate.

### *In vitro* oscillatory pressure model of barotrauma

Oscillatory pressure was applied to primary pneumocyte-AM co-culture at an air-liquid interface using a custom design apparatus as described previously^47^. Briefly, primary pneumocytes and AMs were co-cultured onto Transwell 6-well inserts as described above. Media was removed from the apical chamber and basal chamber media was replaced with fresh HEC basal media (with 1% anti-anti). The well was sealed using rubber stoppers fixed to a central channel connected to the tubing. Tubes from each well were connected to a water manometer and a small animal ventilator (Harvard Apparatus, Holliston, MA, USA). Oscillatory pressure was applied for 16 h with a 0–20 cm-H_2_O range at 0.2 Hz. The media were subsequently collected for analysis by ELISA, and cellular RNA was isolated for RT-qPCR analysis.

### Fluorescence-activated cell sorting

Cells were live/dead stained with Fixable Aqua Dead Cell Stain Kit (Molecular Probes, Eugene, OR, USA) on ice for 30 minutes and protected from light. Cells were blocked with Human TruStain FcX (BioLegend, San Diego, CA, USA) for 20 minutes and stained for 20 minutes with fluorescently labeled antibodies: Alexa Fluor 700 anti-human CD45 (Clone 2D1) and Alexa Fluor 488 anti-human CD326 (clone 9C4), all purchased from BioLegend. Cells were then washed twice in Cell Staining Buffer (BioLegend) and were sorted on a BD FACS Aria-II flow cytometer (BD Biosciences). The data were analyzed with FlowJo software (TreeStar). The flow cytometric gating strategy is presented in **Figure S4**.

### *In vivo* nanoparticle distribution study

*In vivo* distribution was assessed using immunofluorescence staining as described previously^6^. 0.1 nmol Cy3-Dye-labeled Pre-miR^TM^ negative control (Ambion, AM17120) loaded MLNPs or LNPs were administered intratracheally (15uL) into WT mice. Control mice were treated with an equivalent volume of sterile saline without nanoparticles. Mice were euthanized 4 h after nanoparticle administration and the right heart was perfused PBS (Gibco) followed by 2% (v/v) paraformaldehyde (PFA) (ThermoFisher Scientific) in PBS. Lungs were inflated to 20 cm H_2_O and then immersed in 2% PFA for 2 h. After fixation, lung tissues were embedded in optimal cutting temperature (OCT) compound (Fisher Healthcare) and frozen in 2-methylbutane (Honeywell, Charlotte, NC, USA) in a beaker submerged in liquid nitrogen. OCT-embedded lungs were cryosectioned in the coronal plane at 7 μm using a cryostat microtome (Leica CM 1510S). Cryosections were fixed on slides with 2% PFA for 15min at room temperature and washed three times with 1X PBS. Cryosections were permeabilized with 0.1% (v/v) TritonX-100 (Sigma) in PBS for 5min, then incubated in blocking solution, 1% bovine serum albumin (BSA) (Fisher Scientific) in PBS, for 1 h at room temperature. The sections were immunostained for CD68 to identify macrophages using fluorescently conjugated primary antibodies at a 1:50 dilution overnight at 4 °C. Rat anti-CD68 antibody-Alexa Fluor 488 (ab201844) was purchased from Abcam. Sections were washed three times with 1X PBS. Cryosections and coverslips were mounted with Immu-mount (Thermo Scientific). Images were acquired with a fluorescence microscope (Olympus IX81-ZDC) and processed using Fiji software. Nanoparticle distribution was quantitated by co-localization analysis of fluorescence images using Image Calculator in Fiji software. Nanoparticle distribution in macrophages was measured as pixel area overlapping between green (macrophage-Alexa Fluor 488) and red (Cy3 nanoparticle) channels. The percentage co-localization was calculated as the ratio of macrophage cell pixel area overlapping with red nanoparticle pixel area to the total pixel area of macrophages. The percentage of co-occurrence was quantified by manual counting of the number of macrophage cells containing nanoparticles by a second investigator who was blinded to treatment groups.

### Mouse model of hemorrhagic shock-induced ARDS (HS-ARDS)

Prior to the experiment, a pressure catheter with tip size 1F (1/3mm) (Millar Inc., Houston, TX, USA) was soaked in sodium chloride (NaCl) for at least 30 minutes. The pressure catheter was then connected to a Power Lab System and calibrated according to the manufacturer’s protocol. Male wild-type mice were anesthetized with 10 μg ketamine and 1 μg xylazine per gram of animal body weight. Fur was removed from the bilateral inguinal regions using a depilatory cream. A 4-5 mm incision was made in the skin parallel to the left internal oblique muscle and left transverse abdominal muscle. The adipose tissue was separated using fine-tipped forceps to expose the abdominal connection. The adipose tissue and fascia were teased away using Dumont forceps. The femoral triangle (vein, artery, and nerve) was identified below the adipose tissue, at the abdominal connection, and then the nerve was dissected away by grabbing the adjacent adipose tissue. A total of three size 8-0 silk sutures (Fine Science Tools, Foster City, CA, USA) were placed around the vein and artery. The distal artery was ligated and a small arteriotomy incision was made at the proximal portion using micro scissors. The incision was opened using the Dumont forceps by carefully placing one end of the Dumont into the arterial vessel lumen and closing them over the vessel wall. A pressure catheter was inserted into the lumen of one artery while holding the arterial wall and pulling the vessel over the pressure catheter and secured using a silk suture. The proximal hemostat was then released, and the pressure catheter was gently advanced 4-5 mm into the vessel. The pressure catheter was connected to the Power Lab System for blood pressure monitoring. The contralateral artery was cannulated with a polyethylene catheter at 0.28 mm inner diameter x 0.61 mm outer diameter using a similar technique. This catheter was connected to a 3-way stopcock and a 1 mL syringe containing 0.2 mL heparinized sterile saline. Blood was withdrawn into the catheter and mixed with the heparinized saline to prevent clotting. Blood was withdrawn slowly over about 10 minutes until the mean arterial blood pressure (MAP) reached 30-40 mmHg. The BP was maintained in this range for 30 minutes. The mouse was then resuscitated over a 15 min interval by transfusing the approximately 3 times shed blood and lactated ringers’ solution until the mean arterial BP reached 70-90 mmHg. The catheters were removed, and the wounds were closed with sutures.

Following the hemorrhagic shock, mice were treated with 0.1 nmol miR-146a loaded MLNPs or LNPs by intra-tracheal injection (15uL). Mice administered with 0.1 nmol miR-scramble loaded MLNPs were used as controls. A tracheostomy cannula was placed, and mice were connected to the Flexivent small animal ventilator (Scireq, Montreal, QC, Canada). Mice’s body temperature was maintained by using a homeothermic monitoring system (Harvard Apparatus, Holliston, MA, USA). Animals were volume resuscitated with 300-500 μl sterile saline by intraperitoneal (IP) injection. Mice were ventilated for 2 h using a tidal volume (TV) of 12 ml/kg to induce volutrauma and 0 cm-H_2_O PEEP to induce atelectrauma using the ventilator (Scireq). Blood oxygen saturations were measured using a mouse thigh pulse oximeter (Starr Life Sciences). Baseline lung physiology parameters (*e.g*., lung elastance, airway resistance) were obtained by performing a recruitment maneuver followed by forced oscillation^5^. At each 30 minutes time-point, the lungs were recruited (two deep inflations) and lung physiology parameters were measured. At the end of the ventilation, mice were sacrificed by injecting a lethal overdose of anesthetic. BAL was performed by instilling 1 ml of saline into the lungs twice and withdrawing it each time, repeated twice. BAL fluid was then centrifuged for 10 min at 2000 rpm at 4°C and the supernatant was collected for subsequent analysis. The pelleted cells were lysed with RBC lysis buffer (Alfa Aesar, Haverhill, MA, USA). BAL cells were counted with an Automated Cell Counter (Bio-Rad), and differential staining was performed with a Hema 3 Stat Pack (Thermofisher Scientific) according to the manufacturer’s protocol. The remaining cells were pelleted and resuspended in TRIzol (Ambion) for RNA extraction. Following BAL, the left lung was inflated and fixed with 10% formalin. Other organs including the right lungs, liver, heart, spleen, kidney, and diaphragm were harvested and snap-frozen in liquid nitrogen. For whole lung gene expression, the right lungs were placed into a tube containing 600 μl of TRIzol (Ambion), homogenized using a handheld tissue homogenizer (Omni International, Kennesaw GA, USA) purified as described above.

### ELISA

Enzyme-linked immunosorbent assay (ELISA) was performed to evaluate secreted cytokine/mediator levels in cell culture media or BAL according to the manufacturer’s protocol. For human cytokines (IL6, IL8, MIP-1α/β),, OptEIA kits (BD Biosciences, Franklin Lakes, NJ, USA) were used (IL6, IL8, MIP-1α/β) and the manufacturer’s protocol was followed. For mouse mediators (IL6, CXCL1/KC), Duoset ELISA kits (R&D Systems, Minneapolis, MN, USA) were used (IL6, CXCL1/KC) and the manufacturer’s protocol was followed.

### Immunoblotting

Primary human alveolar macrophages (AMs) were lysed in RIPA buffer (VWR, Radnor, PA, USA) supplemented with protease (Roche Diagnostics GmbH, Mannheim, Germany) and phosphatase inhibitors (Sigma Aldrich). Cells were disrupted by a freeze/thaw cycle. Protein extracts were centrifuged to pellet insoluble debris, protein concentration was measured by BCA assay (Thermo Fisher Scientific), and protein concentrations were equalized prior to boiling in the recommended sample buffers supplemented with 2.5% beta-mercaptoethanol (Bio-Rad). SDS-PAGE electrophoresis was performed using 4–12% gradient bis-tris gels (Invitrogen), and proteins were transferred to nitrocellulose membranes (Bio-Rad). Membranes were blocked in 5% w/v nonfat dry milk in tris-buffered saline with 0.1% Tween 20 (TBST) for 1 h at room temperature, and then incubated in primary anti-mannose receptor (CD206) antibody (Abcam, Cambridge, UK) overnight at 4°C. After incubation with HRP-conjugated secondary antibodies, proteins were visualized using SignalFire Elite ECL Reagent (Cell Signaling Technology, Danvers, MA, USA) and imaged using a ChemiDoc XRS+ System (Bio-Rad).

### Human Cytokine Array

The supernatant was collected following the centrifugation of cell culture media. Cytokines were detected using a human cytokine array C1000 kit (RayBiotech, Norcross, GA, USA) according to the manufacturer’s protocol. Briefly, supernatant samples were added onto the antibody membranes and incubated overnight at 4°C following 30 min blocking with blocking buffer. The membranes were incubated with a biotinylated antibody cocktail for 2 h at room temperature and then incubated with HRP-Streptavidin for another 2 h at room temperature. Membranes were transferred onto a plastic sheet, detection buffer mixture was applied onto the membrane and incubated for 2 h at room temperature. The membrane was covered with another plastic sheet and immediately imaged using a ChemiDoc XRS+ System (Bio-Rad).

### Statistics

Statistical analysis was performed using Graphpad Prism 9. Outliers were identified using the ROUT method with a Q threshold of 1%. The distribution of all data was tested for normality and lognormality *via* a Shapiro-Wilk test. All data are presented as Min to Max Show all points, except where noted otherwise. For comparisons of 2 groups that were not normally distributed, a Mann–Whitney test was used. For comparison among multiple groups, for data that were normally or log-normally distributed, an ANOVA was performed on untransformed or log-transformed data as appropriate; for data that were not normally or log-normally distributed, a nonparametric Kruskal-Wallis test was performed. For experiments with two independent variables, a two-way ANOVA was performed after testing data for normality as described. If a significant effect was determined by ANOVA, a Tukey or Sidak post-hoc test for individual group comparisons was performed. A p-value of <0.05 was considered statistically significant.

## ASSOCIATED CONTENT

### Supporting Information

Detailed supplementary results for mannose receptor expression on primary alveolar macrophages, encapsulation efficiency of miR-146a loaded MLNPs, cellular uptake of miR-146a loaded MLNPs in primary AMs, representative flow cytometric gating strategy for identification of pneumocytes and primary AMs, *in vitro* barotrauma increases inflammatory cytokines/chemokines level in the co-culture system, hemorrhagic shock following injurious mechanical ventilation-induced acute respiratory distress syndrome in mice, *in vivo* doseresponse of 9.3-MLNP delivery of miR-146a, *in vivo* miR-146a level in RNA from lung homogenate following nanoparticle delivery and HS-ARDS (PDF).

## Author Contributions

Q.F., S.N.G., and J.A.E. conceived and designed the study. Q. F. performed the majority of experiments and analyzed the data under the supervision of S.N.G. and J.A.E.. Additional data were acquired by E.M.S., R.B.. Data were interpreted by Q.F., V.C.S., M.N.B., L.F., R.L., S.N.G., and J.A.E. The manuscript was drafted by Q.F., S.N.G., and J.A.E. All authors have given approval to the final version of the manuscript.

## Notes

The authors declare no competing interests.

## ACKNOWLEDGEMENTS

This work is partially supported by an R01 HL142767 (JAE, SNG), a Department of Defense Grant W81XWH-19-1-0210 (SNG, JAE), and the Eli Lilly Fellowship in Pharmaceutics at Ohio State (QF). We would like to thank the Ohio State Center for Electron Microscopy and Analysis and Analytical Flow Cytometry Core for their technical assistance. We thank the Lifeline of Ohio Organ Procurement Organization for providing the explanted donor organs, as well as the generous donors and their families whose selfless donations make lifesaving research possible. Specimens and/or data were provided by The Ohio State University Wexner Medical Center DHLRI CTC Human Tissue Biorepository.

