## Supplemental Data for "Macrophage-targeted lipid nanoparticle delivery of microRNA-146a to mitigate hemorrhagic shock-induced acute respiratory distress syndrome"

**Supplementary Figures**

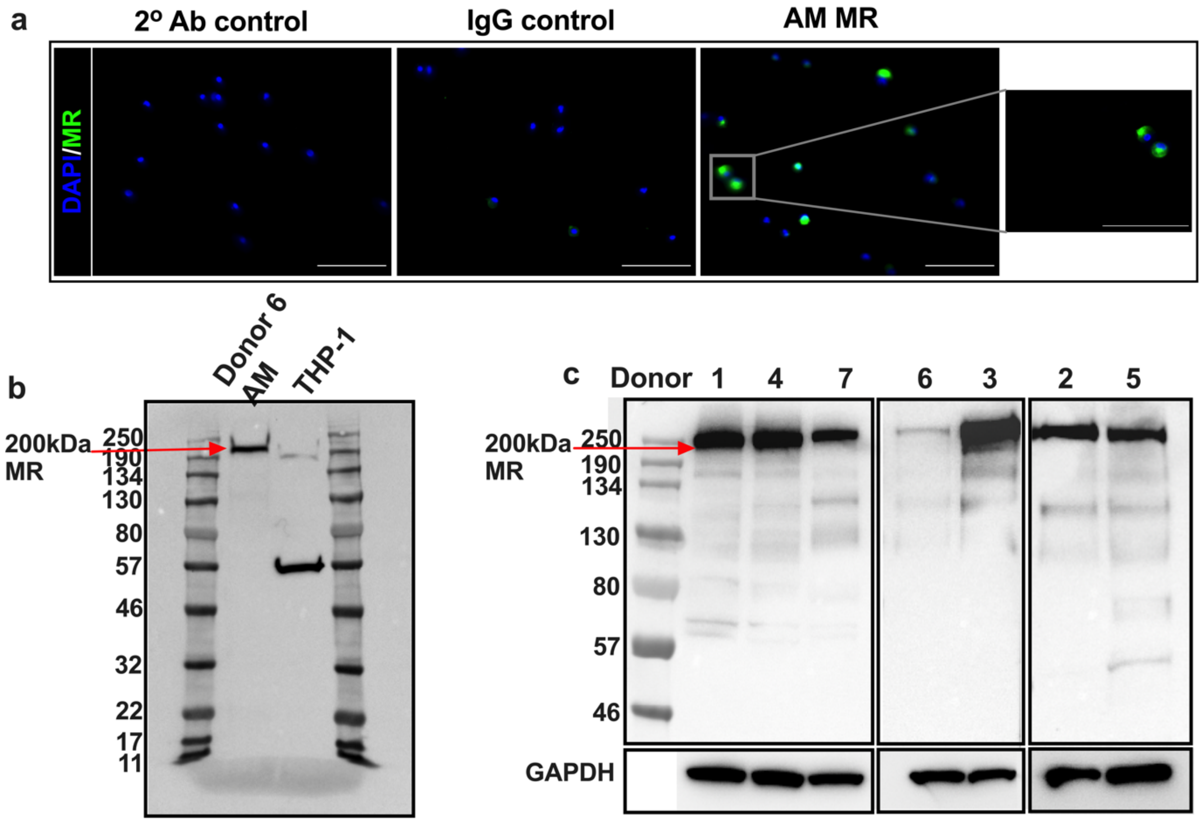


**Figure S1. Mannose receptor expression on primary alveolar macrophages. a)** Representative fluorescence images of primary alveolar macrophages (AMs) immunofluorescence stained for mannose receptor (MR) (CD206) (green) and nuclei (blue). Scale bar: 100 µm. **b)** Representative MR expression assessed by western blotting of AMs and THP-1 cells. **c)** MR expression by western blotting of AMs from 7 donor lungs.

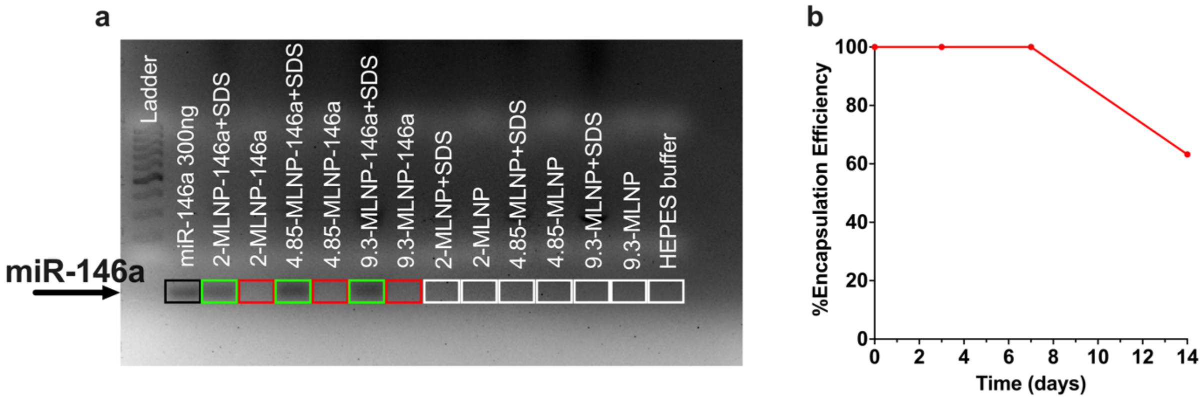

**Figure S2. Encapsulation efficiency of miR-146a loaded MLNPs. a)** Agarose gel electrophoresis demonstrating loading of miR-146a into various MLNP formulations. Boxes show bands for analysis where green box indicates miR levels measured when MLNPs were treated with sodium dodecyl sulfate (SDS) and red box indicates miR levels for intact MLNPs. Black box is the free miR positive control and white boxes are the empty MLNPs and HEPES buffer controls. **b)** Percentage of encapsulation efficiency of miR-146a in MLNP determined by Qubit™ microRNA Assay at time intervals of day 0, 3, 7, and 14.

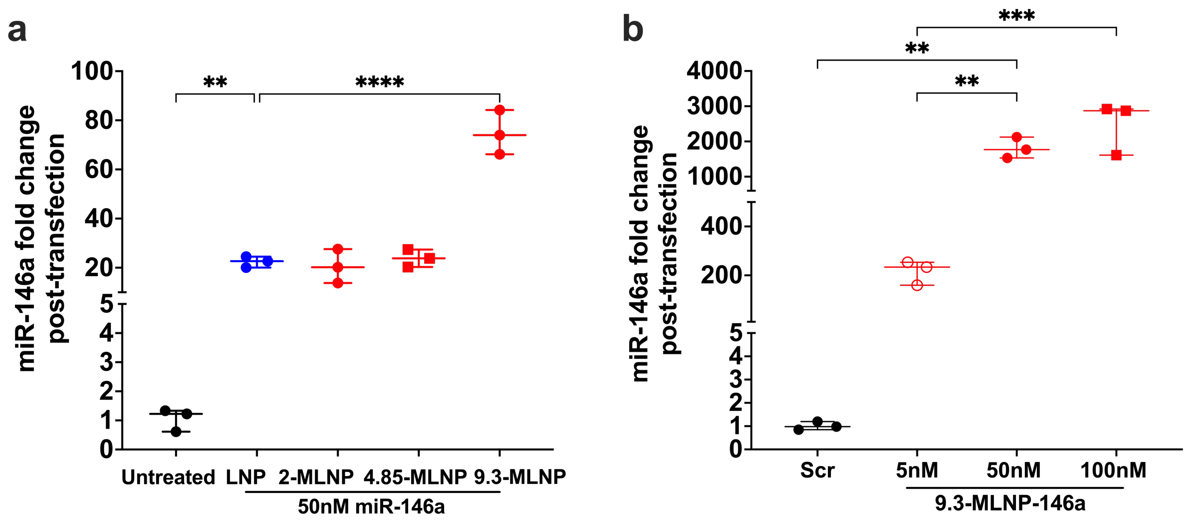


**Figure S3. Cellular uptake of miR-146a loaded MLNPs in primary AMs. a)** Transfection efficiency in term of miR-146a level in primary alveolar macrophages (AMs, donor 6) following delivery of miR-146a with different lipid nanoparticle formulations (non-mannosylated LNPs, mannosylated LNPs containing 2%, 4.85% or 9.3% of mannose-conjugated PA-PEG lipid), calculated by $\Delta\Delta$Ct method, normalized to scramble controls. **b)** Dose-dependent miR-146a level in AMs (donor 4) following 9.3-MLNP delivery of 5nM, 50nM or 100nM miR-146a. Data are normally distributed by Shapiro-Wilk test. Statistical analysis was performed by one-way ANOVA with Tukey's multiple comparisons test. Statistical differences are denoted as **$p<0.01$, ***$p<0.001$, ****$p<0.0001$. Data are presented as Min to Max. Show all points, n=3 wells per group.

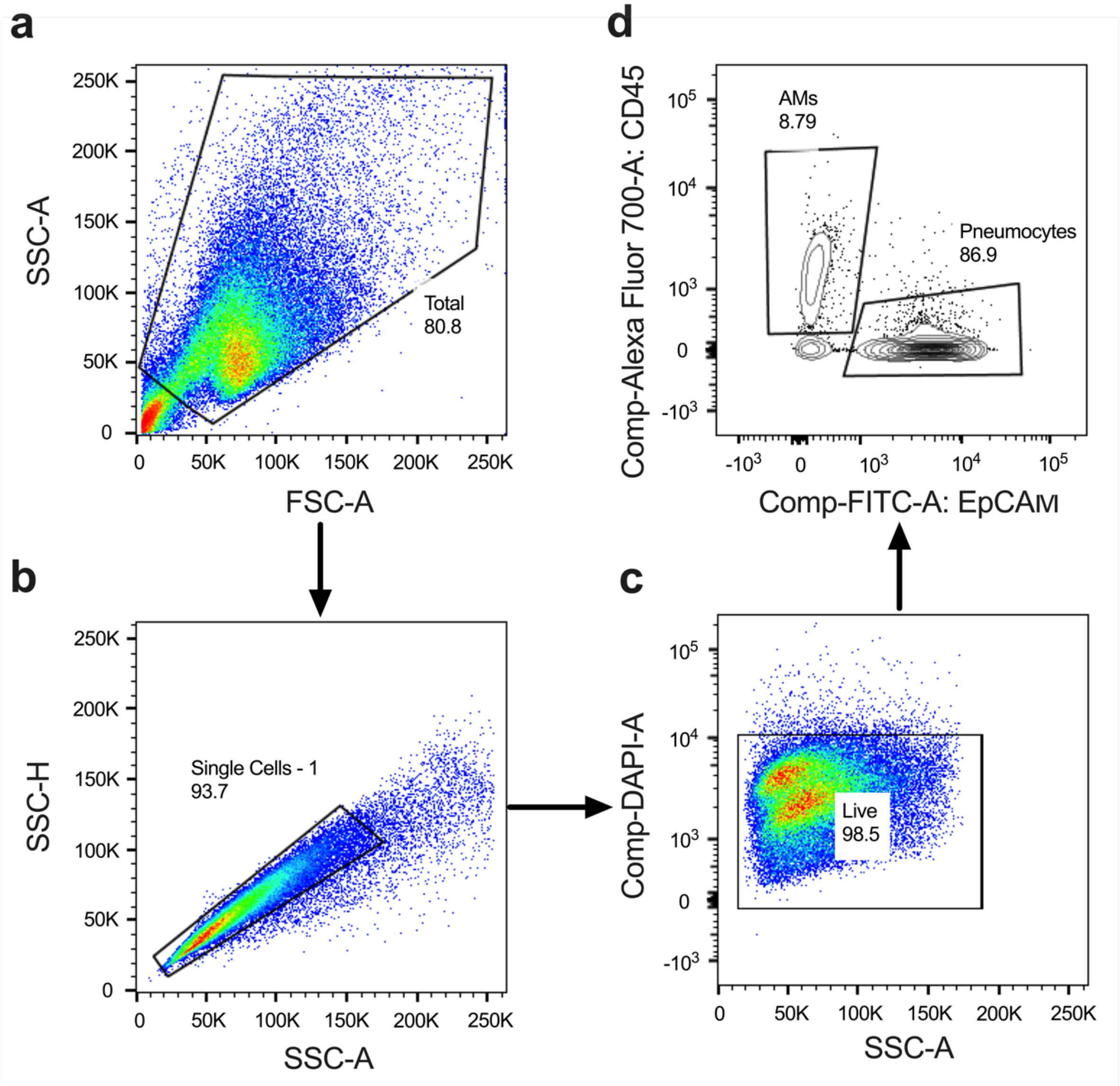


**Figure S4. Representative flow cytometric gating strategy for identification of pneumocytes (EpCAM+) and primary AMs (CD45+).** Total cells were first gated on a forward scatter (FSC) and side scatter (SSC) plot **a)** to exclude debris and then gated to select single cells **b)**. The single cell population was then gated on live cell population **c)**. These live cells were further gated for EpCAM+ cells and CD45+ cells **d)**. Data were processed using FlowJo software.

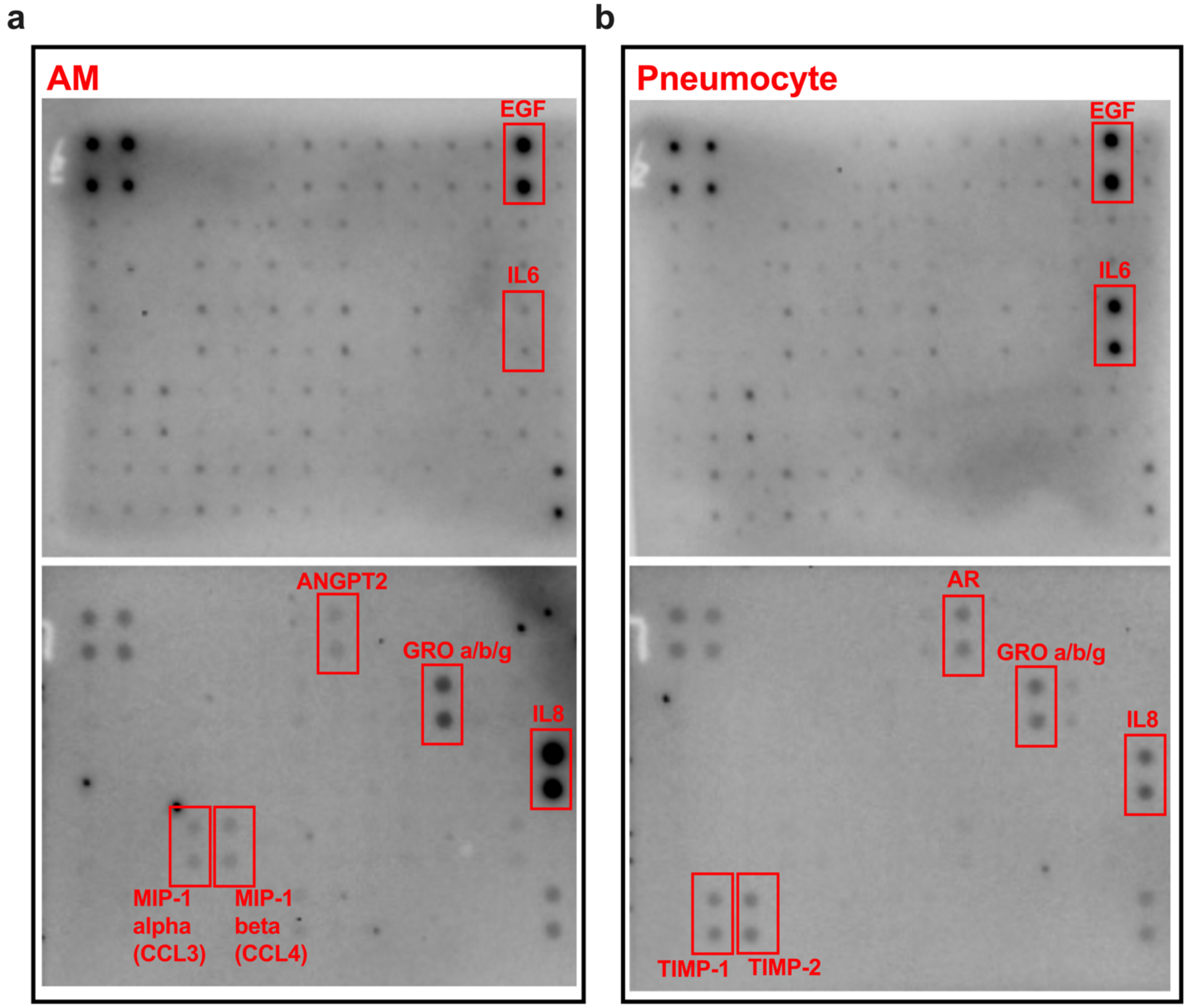


**Figure S5.** ***In vitro*** **barotrauma increases levels of inflammatory cytokines/chemokines in co-culture system.** Human cytokine/chemokine array detection of supernatant from **a)** AMs (donor 5) and **b)** pneumocytes subjected to 16 h in vitro barotrauma (oscillatory pressure) at an air-liquid interface.

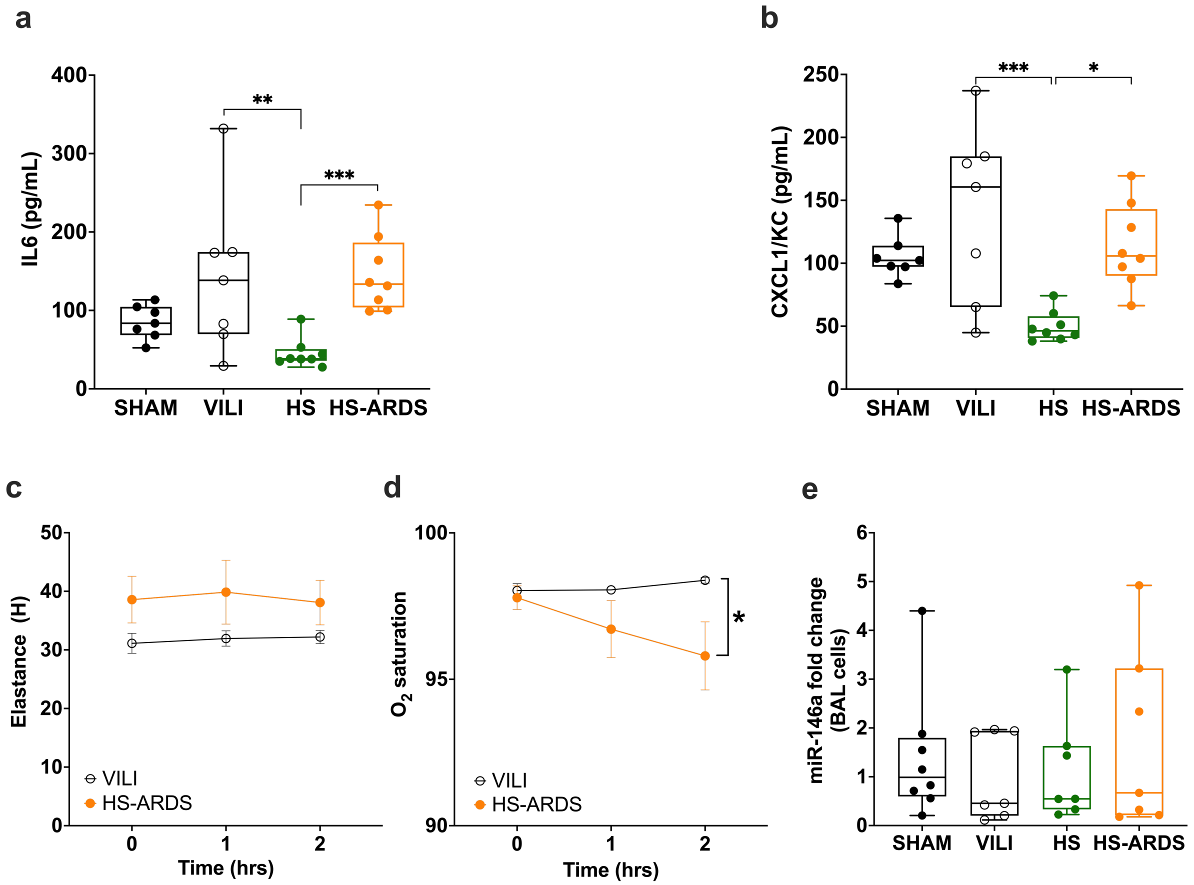


**Figure S6. Hemorrhagic shock following injurious mechanical ventilation induces acute respiratory distress syndrome in mice. a)** Bronchoalveolar lavage (BAL) IL6 concentrations from Sham, ventilator-induced lung injury (VILI), hemorrhagic shock (HS), hemorrhagic shock-induced ARDS (HS-ARDS); Data are lognormally distributed by Shapiro-Wilk test. *p=0.0024, **p = 0.0002 via one-way ANOVA with Tucky's multiple comparisons test, n = 8 per group. **b)** CXCL1/KC concentrations from Sham, VILI, HS and HS-ARDS groups. *p=0.0151, ***p = 0.0008 via one-way ANOVA with Tucky's multiple comparisons test, n = 8 per group. **c)** Lung tissue elastance measurements at initiation (baseline, 0 h), 1 h and at the conclusion of 2 h ventilation. Data are normally distributed by Shapiro-Wilk test. Nonsignificant via two-way ANOVA with Sidak's multiple comparison test, n = 8 per group. **d)** Blood oxygen saturation (SpO2) was measured by pulse oximetry at initiation, 1 h and conclusion of ventilation. Data are normally distributed by Shapiro-Wilk test. *p=0.0226 via two-way ANOVA with Sidak's multiple comparison test, n = 8 per group. **e)** miR-146a fold change from RNA extracted from BAL cells following VILI, HS, HS-ARDS or from Sham controls. Relative expression determined with ΔΔCt method, normalized to Sham. Data are lognormally distributed by Shapiro-Wilk test. Nonsignificant via one-way ANOVA with Tukey's multiple comparison test, n = 8 per group. Data are presented as Min to Max. Show all points.

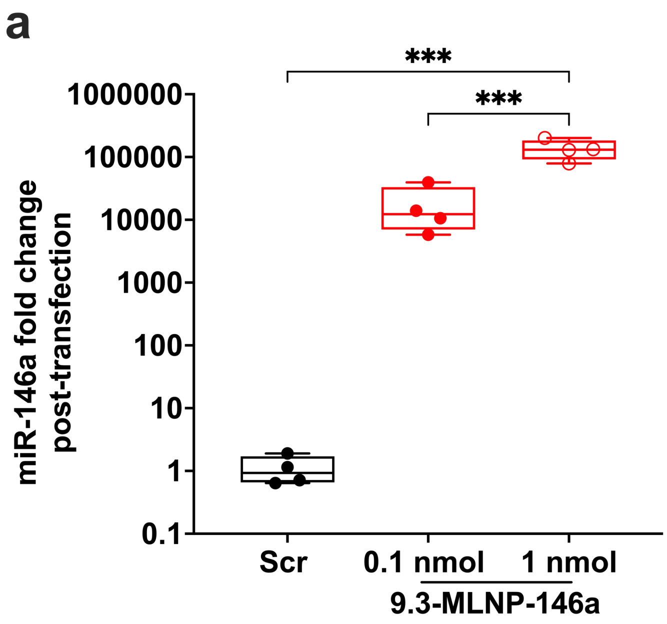


**Figure S7. *In vivo* dose-response study of 9.3-MLNP delivery of miR-146a. a)** Dose-dependent miR-146a level in BAL cells following 9.3-MLNP delivery of 0.1 nmol, or 1 nmol miR-146a. Data are normally distributed by Shapiro-Wilk test. ***p<0.001 via one-way ANOVA with Tukey's multiple comparisons test, n=4 per group. Data are presented as Min to Max. Show all points.

a

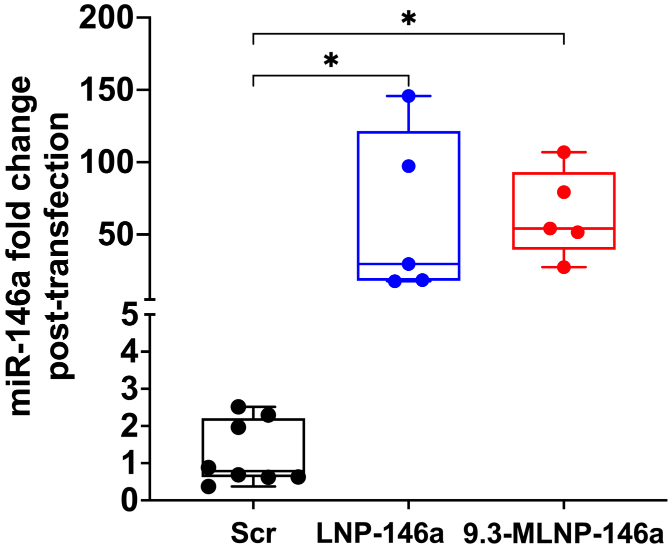


**Figure S8.** ***In vivo*** **miR-146a level in RNA from lung homogenate following nanoparticle delivery in HS-ARDS mice.** Relative expression determined by ΔΔCt method, normalized to scramble control. Data are normally distributed by Shapiro-Wilk test. *$p<0.05$ via one-way ANOVA with Tukey's multiple comparison test. n=8 for scramble group and n=6 for miR groups.
